# Bayesian Inference Of Phylogenetic Networks From Bi-allelic Genetic Markers

**DOI:** 10.1101/143545

**Authors:** Jiafan Zhu, Dingqiao Wen, Yun Yu, Heidi M. Meudt, Luay Nakhleh

**Affiliations:** Computer Science, Rice University, Houston, TX, USA; Museum of New Zealand Te Papa Tongarewa, Wellington, New Zealand; BioSciences, Rice University, Houston, TX, USA

## Abstract

Phylogenetic networks are rooted, directed, acyclic graphs that model reticulate evolutionary histories. Recently, statistical methods were devised for inferring such networks from either gene tree estimates or the sequence alignments of multiple unlinked loci. Bi-allelic markers, most notably single nucleotide polymorphisms (SNPs) and amplified fragment length polymorphisms (AFLPs), provide a powerful source of genome-wide data. In a recent paper, a method called SNAPP was introduced for statistical inference of species trees from unlinked bi-allelic markers. The generative process assumed by the method combined both a model of evolution for the bi-allelic markers, as well as the multispecies coalescent. A novel component of the method was a polynomial-time algorithm for exact computation of the likelihood of a fixed species tree via integration over all possible gene trees for a given marker. Here we report on a method for Bayesian inference of phylogenetic networks from bi-allelic markers. Our method significantly extends the algorithm for exact computation of phylogenetic network likelihood via integration over all possible gene trees. Unlike the case of species trees, the algorithm is no longer polynomial-time on all instances of phylogenetic networks. Furthermore, the method utilizes a reversible-jump MCMC technique to sample the posterior of phylogenetic networks given bi-allelic marker data. Our method has a very good performance in terms of accuracy and robustness as we demonstrate on simulated data, as well as a data set of multiple New Zealand species of the plant genus *Ourisia*(Plantaginaceae). We implemented the method in the publicly available, open-source PhyloNet software package.

**Author summary:** The availability of genomic data has revolutionized the study of evolutionary histories and phylogeny inference. Inferring evolutionary histories from genomic data requires, in most cases, accounting for the fact that different genomic regions could have evolutionary histories that differ from each other as well as from that of the species from which the genomes were sampled. In this paper, we introduce a method for inferring evolutionary histories while accounting for two processes that could give rise to such differences across the genomes, namely incomplete lineage sorting and hybridization. We introduce a novel algorithm for computing the likelihood of phylogenetic networks from bi-allelic genetic markers and use it in a Bayesian inference method. Analyses of synthetic and empirical data sets show a very good performance of the method in terms of the estimates it obtains.

## Introduction

The availability of genome-wide data from many species and, in some cases, many individuals per species, has transformed the study of evolutionary histories, and given rise to phylogenomics—the inference of gene and species evolutionary histories from genome-wide data. Consider a data set *S* = {*S*_1_,…, *S_m_*} consisting of the molecular sequences of *m* loci under the assumptions of free recombination between loci and no recombination within a locus. The likelihood of a species phylogeny Ψ (topology and parameters) is given by

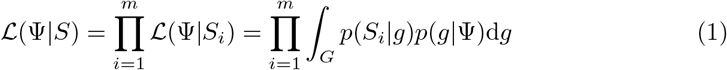

where the integration is taken over all possible gene trees. The term *p*(*S_i_*|*g*) is the likelihood of gene tree *g* given the sequence data of locus *i* [1]. The term *p*(*g*|Ψ) is the density function (pdf) of gene trees given the species phylogeny and its parameters. For example, Rannala and Yang [2] derived this pdf under the multispecies coalescent (MSC). This formulation underlies the Bayesian inference methods of [2–4].

Debate has recently ensued regarding the size of genomic regions that would be recombination-free (or almost recombination-free) and could truly have a single underlying evolutionary tree [5,6]. One way to overcome this issue is to use unlinked single nucleotide polymorphisms (SNPs) or amplified fragment length polymorphisms (AFLPs). Such data provide a powerful signal for inferring species phylogenies and the issue of recombination within a locus becomes irrelevant. Furthermore, as long as those markers are sampled far enough from each other the assumption of free recombination among loci holds. Indeed, this is the basis of the SNAPP method that was recently introduced in [7]. Since a bi-allelic SNP or AFLP marker has no signal by itself to resolve much of the branching patterns of a gene genealogy, a major contribution of Bryant *et al*. was an algorithm for analytically computing the integration in Eq. (1) for bi-allelic markers.

While trees constitute an appropriate model of the evolutionary histories of many groups of species, it is well known that other groups of species have evolutionary histories that are reticulate [8]. Horizontal gene transfer is ubiquitous in prokaryotes [9,10], and several bodies of work are pointing to much larger extent and role of hybridization in eukaryotic evolution than once thought [8, 11–15]. Not only does hybridization play an important role in the genomic diversification of several eukaryotic groups, but increasing evidence is pointing to the adaptive role it has played, for example, in wild sunflowers [16], humans [17], macaques [18], mice [19], butterflies [20], and mosquitoes [21, 22].

Reticulate evolutionary histories are best modeled by *phylogenetic networks*. Two statistical methods were recently introduced for inference under the formulation given by Eq. (1), when Ψ is a phylogenetic network [23,24], and other methods were also introduced for statistical inference of phylogenetic networks using gene tree estimates as the input data [25–29].

The methods of [23, 24] assume that the data for each locus consists of a sequence alignment that has no recombination. In this paper, we devise an algorithm that builds on the algorithm of [7] for analytically computing the integral in Eq. (1) when Ψ is a phylogenetic network. In other words, our algorithm allows for computing the likelihood of a phylogenetic network from unlinked bi-allelic markers while analytically integrating out the gene trees for the individual markers. We couple this likelihood function with priors on the phylogenetic network and its parameters to obtain a Bayesian formulation, and then employ the reversible-jump MCMC (RJMCMC) kernel from [23] to sample the posterior of the phylogenetic networks and their associated parameters given the bi-allelic data.

We implemented our algorithm and the RJMCMC sampler in PhyloNet [30], which is a publicly available open-source software package for inferring and analyzing reticulate evolutionary histories. We studied the performance of our method on simulated and biological data. For simulations, we extended the framework of [7] so that the evolution of bi-allelic markers could be simulated within the branches of a phylogenetic network. For the biological data, we analyzed two data sets of multiple New Zealand species of the plant genus *Ourisia* (Plantaginaceae). The results on the simulated data show very good accuracy and robustness as reflected by the method’s ability to recover the true phylogenetic networks and their associated parameters even when the underlying assumptions of the method are violated. For the biological data, the method recovers two established hybrids and their putative parents correctly.

The proposed method and Bayesian sampler provide a new tool for biologists to infer reticulate evolutionary histories, while also account for the complexity arising from incomplete lineage sorting, from bi-allelic markers, thus complementing existing tools that use gene tree estimates or sequence alignments of the individual loci as the input data. The use of such bi-allelic markers, particularly when they are sampled far enough across the genome, completely sidesteps potential problems that could arise due to the presence of recombination within loci.

## Materials and methods

### Phylogenetic networks and gene trees

A *phylogenetic* 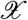-*network*, or 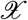-network for short, Ψ is a rooted, directed, acyclic graph (DAG) whose leaves are bijectively labeled by set 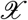 of taxa. We denote by *V*(Ψ) and *E*(Ψ) the sets of nodes and edges, respectively, of the phylogenetic network Ψ. Every node of the network has in-degree 1, which we call a *tree node*, or in-degree 2, which we call a *reticulation node*. The only exception is special node *s* whose in-degree is 0 and out-degree is 1; the edge (*s*,*r*) defines the branch above the root. The edges whose head is a reticulation node are the *reticulation edges* of the network; all other edges constitute the *tree edges* of the network. Every edge is directed forward in time. We assume all phylogenies considered here (trees and networks) are binary—no node has out-degree higher than 2.

Here, we use the bottom of a branch to refer to the end of the branch that is farther from the root of the network, and use the top of a branch to refer to the end of the branch that is closer to the root. Given that the coalescent views the evolution of alleles backward in time, we say that a lineage enters a branch to mean a lineage that exists at the bottom of that branch. Similarly, we say a lineage exits a branch to mean a lineage that exists at the top of that branch.

Each node in the network has a species divergence time and each edge *b* has an associated population mutation rate *θ_b_* = 4*N_b_μ*. This parameter is typically referred to in the literature as the (rescaled) population size. Given the length *τ* of a branch in units of expected number of mutations per site, the length of that branch in coalescent units is 2*τ*/*θ*, assuming diploid individuals. The branch above the root, (*s*,*r*), is infinite in length so that all lineages that enter it coalesce on it eventually.

For every pair of reticulation edges *e*_1_ and *e*_2_ that share the same reticulation node, we associate an inheritance probability, *γ*, such that *γ_e_*_1_, *γ*_*e*_2__ ∈ [0,1] with *γ*_*e*_1__ + *γ*_*e*_2__ = 1. We denote by Γ the vector of inheritance probabilities corresponding to all the reticulation nodes in the phylogenetic network.

We use Ψ to refer to the topology, species divergence times, population mutation rates, and inheritance probabilities of the phylogenetic network. That is, here we include Γ as part of Ψ.

An 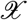-phylogenetic tree, or 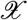-tree, is an 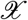-network with no reticulation nodes. A gene tree is an 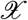-tree. Each node in the gene tree has an associated coalescence time. In the algorithm below, we make use of a coloring function *c*: (*E*(*g*),*t*) → {0,1}, similar to that used in [7], where *c*(*e*,*t*) indicates the color, or allele, at time *t* along the branch *e* of gene tree *g*. We will follow [7] in calling the two colors red and green.

### Labeled partial likelihoods

Looking forward in time (from the root toward the leaves), let *u* and *υ* be the mutation rate from red allele to green allele and the mutation rate from green allele to red allele, respectively. The stationary distribution of the red and green alleles at the root is given by *υ*/(*u* + *υ*) and *u*/(*u* + *υ*), respectively. Observed alleles are indicated by values of the coloring function *c* at gene tree leaves.

Given a gene history embedded within the branches of the network, the numbers and types of lineages at both ends of each branch of the network are needed to compute the likelihood. Let *x* be a branch in the phylogenetic network. We denote by 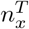 and 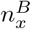 the total numbers of lineages at the top and bottom of *x*, respectively, and by 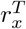 and 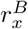 the numbers of red lineages at the top and bottom of *x*, respectively. See Fig. 1 for an illustration.

**Fig 1.**
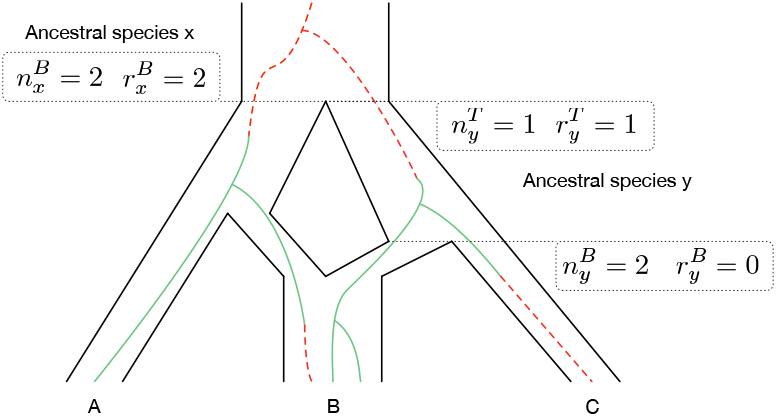
Illustrating the “growth” of lineages of a gene tree in a phylogenetic network. The histories of green and red alleles are shown as solid (green) lines and dashed (red) lines, respectively.

Let *x* be an arbitrary branch in the phylogenetic network and let *R_x_* be the event that for every external branch *z* that is a descendant of *x*, the actual number of red alleles in *z* equals to 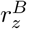.

We define two partial likelihoods: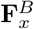 is the product of the likelihood of a subtree rooted at the bottom of *x* and the probability 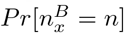, and 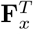 is the product of the likelihood at the top of branch *x* and the probability 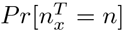. In the case of a species tree (i.e., no reticulation nodes in the species phylogeny), the partial likelihood vectors 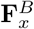 and 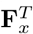 are given by [7]

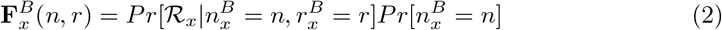

and

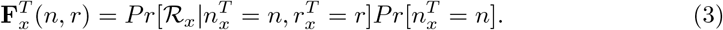

Here 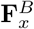 and 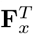 are indexed by nonnegative integers *n* and *r*, where *r* ≤ *n*. Let *M* be the maximum possible value of 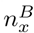 and 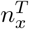 over all branches. Then, each of 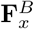 and 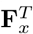 has at most *l* = (1 + (*M* + 1))(*M* + 1)/2 entries.

In the case of a species tree, the path from a leaf to the root is unique. However, this might not be the case for phylogenetic networks: If there is a reticulation node on a path from a leaf to the root, then multiple paths exist between that leaf and the root. This is the issue that necessitates modifying the algorithm of [7] significantly, and that leads to much larger computational requirements in the case of phylogenetic networks. The key idea behind the modification is as follows. As the algorithm proceeds to compute the likelihood in a bottom-up fashion from the leaves to the root, whenever a reticulation node is encountered, the current set of lineages is bipartitioned in every possible way so that one side of the bipartition tracks one parent of the reticulation node and the other side tracks the other parent. As the network has a unique root, the two sides of each bipartition eventually come back together at an ancestral node. At that point, these two sides are merged properly.

To achieve this proper merger, we introduce “labeled partial likelihoods,” or LPL. Like the case of [7], LPLs are not “real” partial likelihoods. The reason for this is that when partial likelihood vectors are split (described below), those become symbolic terms that do not evaluate to partial likelihoods until they are merged later. This is analogous to the difference between *ancestral configurations* on species trees [31] and their labeled counterparts on phylogenetic networks [32], where the latter are in many cases just symbolic terms that do not evaluate to true (partial) likelihood values.

Given a phylogenetic network Ψ with *k* reticulation nodes numbered 0,1, …, *k* – 1, an LPL *P* is an element of [0,1]^*l*^ × ℤ^*k*^, where the first element of the pair is a partial likelihood as in [7]. The second element is the label to keep track of partial likelihoods that originated from a split of the same partial likelihood at a reticulation node so that these two could be merged. More formally, we say two LPLs *P*_1_ = (**F**_1_, s_1_) and *P*_2_ = (**F**_2_, s_2_), where |s_1_| = |s_2_|, are compatible if and only if for every 0 ≤ *i* < |s_1_|, either s_1_(*i*) = s_2_(*i*) or s_1_(*i*) · s_2_(*i*) = 0.

We denote by 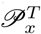 and 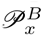 the sets of LPLs that are associated with the top and bottom of branch *x*, respectively. These two quantities are computed in a bottom-up fashion, proceeding from the leaves of the network towards its root. Once the LPLs at the root are computed, the overall likelihood of a given site is computed. As the algorithm proceeds from the leaves towards the root, it needs to compute LPLs at the leaves, the top of a branch, the bottom of reticulation edges, and the bottom of tree edges. We now describe each of those computations; the overall algorithm is simply a bottom-up traversal of the network while applying the appropriate computation as a node is encountered.

### Computing LPLs for leaf nodes

Consider an external branch *x* that is connected to a leaf node. Let *n_x_* denote the number of individuals sampled from the species associated with that leaf, and let *r_x_* be the number of red lineages among those individuals. We create LPL 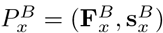, where

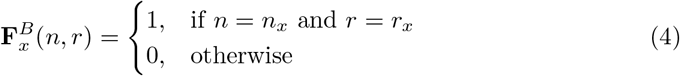

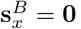. Finally, we associate 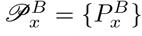 with the bottom of branch *x*.

As pointed out in [7], the input data may contain dominant markers like AFLPs, which means heterozygotes and homozygotes are not distinguishable for the dominant band. If there are dominant markers in the data, and the red allele is dominant, 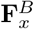 is computed by

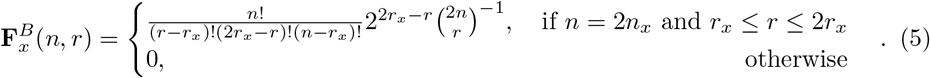

instead of using Eq. (4), where *r* – *r_x_* is the number of homozygous red individuals, 2*r_x_* – *r* is the number of heterozygous individuals (red and green), and *n* – *r_x_* is the number of homozygous green individuals. There are more details about this computation in [7].

### Computing LPLs at the top of a branch

Bryant *et al.* [7] computed partial likelihoods using a continuous-time Markov chain whose transition rate matrix ℚ is indexed by ((*n*, *r*); (*n*′,*r*′)) for transitioning from *n* lineages *r* of which are red alleles to *n*′ individuals *r*′ of which are red alleles, and its entries are given by

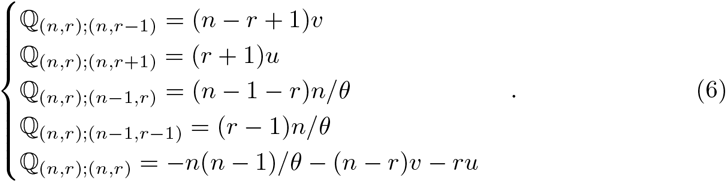

Let *x* be any branch in the phylogenetic network, with *θ* and *t* being the population mutation rate and branch length of *x*, respectively, and assume 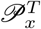 has already been computed. Then,

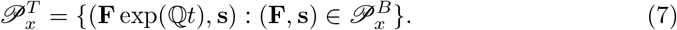

### Computing LPLs at the bottom of reticulation edges

Consider a reticulation node given by two reticulation edges *y* and *z*, with inheritance probabilities *γ* and 1 – *γ*, respectively, and branch *x* emanating from the reticulation node, as illustrated by Fig. 2. The main idea in this part is as follows. Given a set of lineages at the top of branch *x*, a subset of those lineages is inherited along branch *y* and the remaining lineages is inherited along branch *z*. Since there are multiple ways of bipartitioning the set of lineages, the labels in an LPL allow the algorithm to keep track of the subsets of lineages that originated from the same split. We now describe this formally.

**Fig 2.**
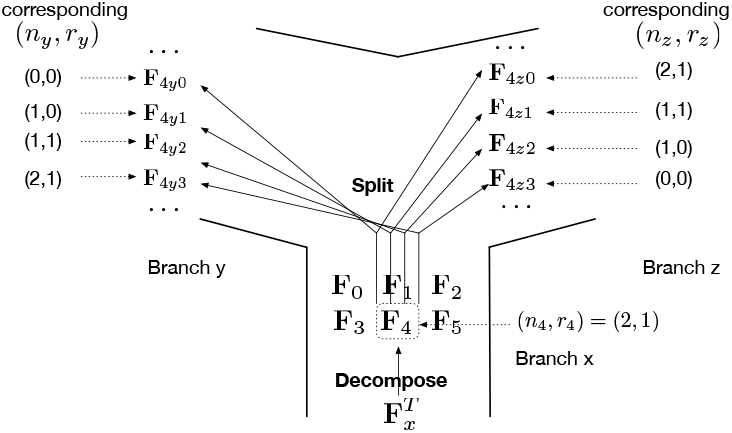
An illustration of the decompose-and-split operation. In this example, partial likelihood 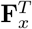 is decomposed into six vectors **F**_0_ to **F**_5_. An illustration of how **F**_4_ is split in the four possible ways to trace branches *y* and *z* is shown, and every split is assigned a unique label.

#### Decomposing

Let (**F**, s) be an LPL in 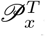. Given that **F** has *l* entries, we decompose **F** into *l* vectors, each with *l* entries: **F**_0_, **F**_1_, …, **F**_*l*-1_. Let *ϕ*: {(*n*′, *r*′): *n*′, *r*′ ∈ ℕ, *r*′ ≤ *n*′ ≤ *m*} → ℕ be given by *ϕ*(*n*′, *r*′) = *n*′(*m* + 1) + *r*′. The entries of **F**_*i*_ are set according to

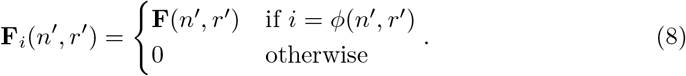

#### Splitting

Consider vector **F**_*i*_ and assume *i* = *ϕ*(*n_i_*,*r_i_*). The existence of *n_i_* lineages out of which *r_i_* are red at the top of branch *x* means that any 0 ≤ *n_y_* ≤ *n_i_* lineages of those could be inherited along branch *y*, and out of those 0 ≤ *r_y_* ≤ min(*r_i_*,*n_y_*) could be red; the remaining *n_z_* = *n_i_* – *n_y_* lineages, out of which *r_z_* = *r_i_* – *r_y_* are red, are inherited along branch *z*. Such a split gives rise to two LPLs: *P_y,n_y_,r_y__* = (**F***_y,n_y_, r_y__*, s*_y,n_y_, r_y__*) and *P_z,n_z_,T_z__* = (**F***_z,n_z_,T_z__*, s*_z,n_z_,r_z__*) with s*_y, n_y_, r_y__* and s_*z, n_z_, r_z_*_ assigned the same value that is unique to the specific split. For this specific split we define

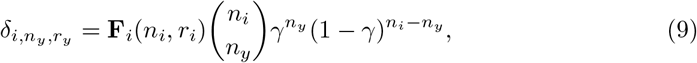

and compute **F**_*y, n_y_, r_y_*_ and **F**_*z,n_z_, r_z_*_ by

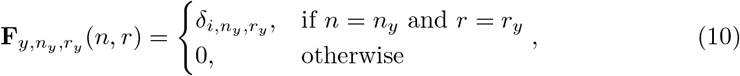

and

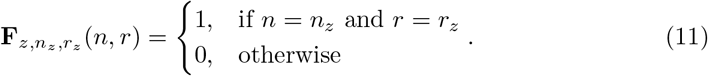

The resulting *P_y_* and *P_z_* from all possible splits constitute the elements of the sets 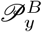 and 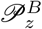, respectively. The full procedure for executing the decompose-and-split operations is given in Algorithm 1.

**Algorithm 1:**
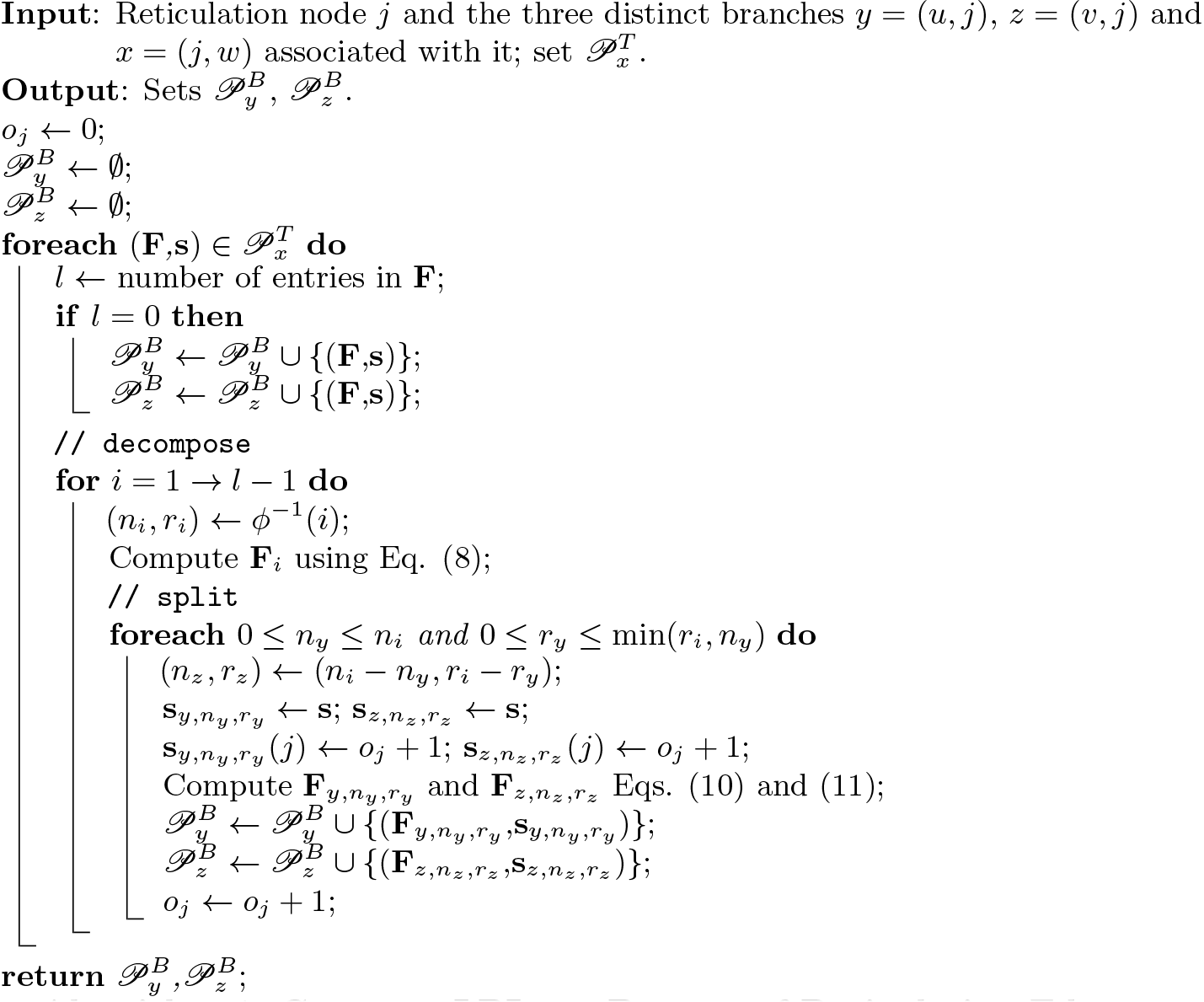
Compute LPLs at Bottom of Reticulation Edges.

Let *n* be the number of individuals in branch *x* and all its descendants. In the middle for loop, *l*, which is the number of entries in each **F**, equals to *O*(*n*^2^). The inner foreach loop over *n_i_* and *r_i_* runs for *O*(*n*^2^) times. Therefore the number of pairs of new LPLs generated is *O*(*n*^4^) for each LPL in branch *x*.

### Computing LPLs at the bottom of a tree edge

Consider an internal tree node *j* with its three associated edges *x* = (*u*,*j*), *y* = (*j*, *υ*), and *z* = (*j*, *w*). We are interested in computing the set 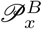 in terms of the two sets 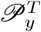 and 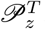. The labels in LPLs allow the algorithm to determine whether two LPLs originated from a split at a descendant reticulation node or not (including the case of no descendant reticulation nodes of node *j*). Let *P_y_* = (**F**_*y*_, s_*y*_) and *P_z_* = (**F**_*z*_, s_*z*_) be two elements of 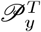 and 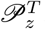, respectively, that are compatible. A label s_*x*_ is computed by

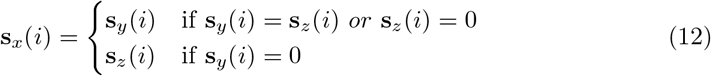

for 0 ≤ *i* ≤ |s_*y*_ |. Furthermore, **F**_*x*_ is computed by

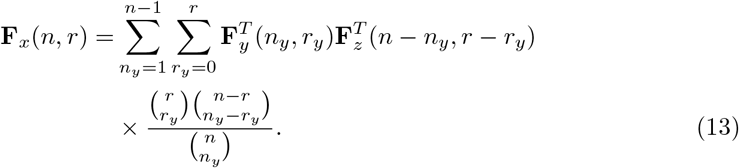

The LPL (**F**_*x*_, s_*x*_) is added to 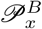. The full procedure for computing set 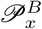 is given in Algorithm 2.

**Algorithm 2:**
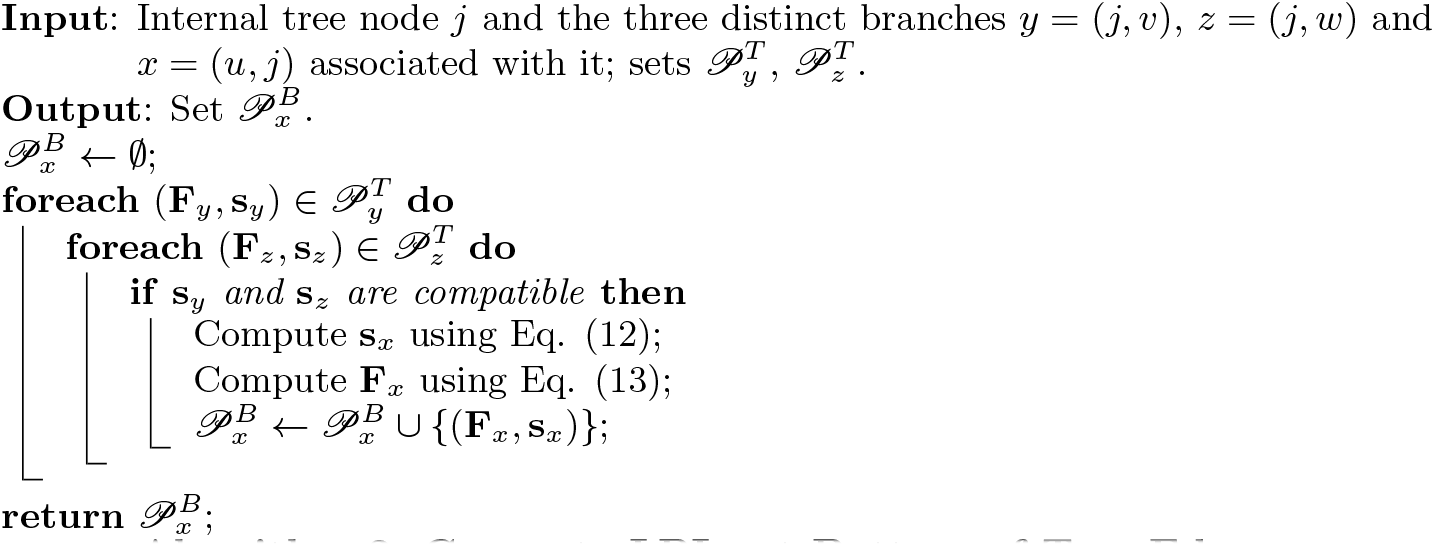
Compute LPLs at Bottom of Tree Edge.

### Termination: Computation above root node

Let the infinite-length branch associated with root be *ρ*. Then, we let 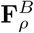 be the sum of all vectors **F** in elements (**F**, s) of set 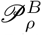.

To obtain the overall likelihood *ℒ*(Ψ|*S_i_*) given the data *S_i_* for site *i*, vector x is obtained as a nonzero solution of ℚx = 0. Like in [7], x is indexed by (*n*, *r*) pairs, such that x(1,0) + x(1,1) = 1. Hence we have x(*n*, *r*) = *Pr*[*R* = *r*|*N* = *n*], where *N* is the number of lineages and *R* is the number of red lineages, if we sample from a single population of constant size with same distribution for the root allele probabilities and allele frequencies as in [7]. The likelihood is given by

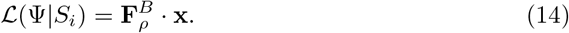

### Optimizing the computation

As described above, the partial likelihood vectors are split to follow every possible way of bipartitioning a set of lineages at a reticulation node. It is this operation that leads to a significant increase in the running time and memory requirement of the likelihood computation as compared to the case of species trees. Here we describe an optimization step that we employ to improve performance in terms of computational requirements, without affecting the correctness of the likelihood computation.

An *articulation node* in a graph is a node whose removal disconnects the graph into two or more components. In a directed graph, a *lowest articulation node* is an articulation node that has at least one child that is neither an articulation node nor a leaf. For example, in a tree, every node is an articulation node. However, in a phylogenetic network that is not necessarily the case. For example, in the phylogenetic network of Fig. 1, the reticulation node is an articulation node. However, the root of the network is the only lowest articulation node (its removal disconnects the special node *s*—the parent of the infinite-length branch above the root—from the rest of the network). The reticulation node is not a lowest articulation node since it has one child and that child is a leaf.

The main idea of the optimization is that all LPLs at a lowest articulation node could be merged into a single LPL, thus avoiding carrying forth all that information. More formally, given a set of LPLs at the bottom of a lowest articulation node, a new LPL is produced by summing all the partial likelihood vectors in the LPLs, and assigning it an empty label. This new LPL is the only one assigned to the bottom of the articulation node; all other LPLs are deleted.

### Time complexity

Our algorithm computes the likelihood of a phylogenetic network given a set of biallelic markers. This algorithm computes matrix exponential along every branch, and processes the network’s nodes in a post-order traversal. Computation at a leaf takes *O*(1) time.

At a reticulation node, the time consumption increases after each reticulation node is processed, due to the accumulation of (split) LPLs. In the last processed reticulation node, the number of LPLs in its descendant is at most *O*(*n*^4(*k*–1)^). There are at most *O*(*n*^4^) new LPLs generated due to decompose-and-split operation for each original LPL. Therefore the time complexity of processing a reticulation node is at most *O*(*n*^4*k*^). We adopted the same approximation of matrix exponential as in [7], so the time complexity of computing matrix exponentiation is *O*(*n*^2^), and computation along every branch is at most *O*(*n*^4*k*+2^).

At a tree node, computation is mostly spent on evaluating Eq. (13). Let *n* be the number of individuals present under an internal tree node. Then, this evaluation takes *O*(*n*^4^) time for a pair of compatible LPLs. The total time consumption of processing tree nodes also depends on the number of LPLs. Assuming *k* reticulation nodes in the phylogenetic network, there are at most *O*(*n*^4*k*^) pairs of compatible LPLs. Therefore the time complexity of processing a tree node is *O*(*n*^4*k*+4^).

In total, the time complexity of the algorithm is *O*(*mn*^4*k*+4^), where *m* is the number of species, *n* is the total number of lineages sampled from the species, and *k* is the number of reticulation nodes. Notice that when *k* = 0, which means the species phylogeny is a tree, the time complexity is *O*(*mn*^4^), which is the running time of the SNAPP algorithm without fast Fourier transforms.

To speed up computation, and since markers are independent, computations for the individual markers are parallelized by multi-threading. Furthermore, the data is preprocessed so that the unique marker patterns are identified and their probabilities are computed only once and reused for for all markers with the same patterns (states for the taxa).

### Bayesian inference

The prior on the phylogenetic network is the same as that employed in [23], which we review briefly here. The prior is given by

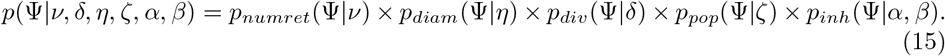

Here, *p*(Ψ|*ν*) is a Poisson prior on the number of reticulation nodes, normalized by the number of networks with the same number of reticulation nodes as Ψ, *p_diam_*(Ψ|*η*) is an exponential prior on the diameters of reticulation nodes. The diameter of a reticulation node is the sum of the branch lengths on the cycle that contains the reticulation node in the underlying undirected graph of the network. *p_div_*(Ψ|*δ*) is an exponential prior on the divergence times. Rannala and Yang used independent Gamma distributions for time intervals (branch lengths) instead of divergence times. However, in the absence of any information on the number of edges of the species network as well as the time intervals, it is computationally intensive to infer the hyperparameters of independent Gamma distributions. Currently, we use a uniform distribution (as in BEST [33]). *p_pop_*(Ψ|*ζ*) is a Gamma prior on the population mutation rate. For *p_pop_*, we use the Gamma distribution Γ(2,*ζ*) with mean value 2*ζ* and shape parameter 2. *p_inh_*(Ψ|*α,β*) is a Beta prior, with parameters *α* and *β*, on the inheritance probabilities. Unless there is some specific knowledge on the inheritance probabilities, a uniform prior on [0, 1] is adopted by setting *α* = *β* = 1.

It is important to note here that if the topology of Ψ does not follow the phylogenetic network definition (e.g., has a cycle), then *p*(Ψ|*ν*, *δ*, *η*, *ψ*) = 0. This is crucial since, in the MCMC kernels we employ for sampling the posterior distribution, we allow the moves to produce directed graphs that slightly deviate from the definition; in this case, having the prior be 0 guarantees that the proposal is rejected. Using the strategy, rather than defining only “legal” moves simplifies the calculation of the Hastings ratios. However, the sampler always guarantees that the divergence times are consistent; that is, no node has a divergence time smaller than or equal to the divergence time of any of its descendants.

We employed the reversible-jump MCMC, or RJMCMC [34] algorithm implemented in PhyloNet [30] to sample from the posterior distribution given by

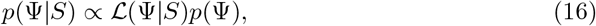

where Ψ here denotes the topology of the network and all its parameters, and *p*(Ψ) is the prior on the network and its parameters as described above.

We make use of only the 12 proposals designed for sampling phylogenetic networks and their parameters described in [23], but not the proposals aimed at sampling gene trees, as gene trees are integrated out.

### Synthetic data generation

We implemented in PhyloNet [30] a program to simulate bi-allelic markers on a given phylogenetic network. Bryant *et al.* [7] simulated bi-allelic markers by first generating gene trees inside a species tree (under the multispecies coalescent model), and then simulating the markers down the gene trees. In our case, we replaced the first step by generating gene trees inside a phylogenetic network under the multispecies network coalescent [26]; the second step of simulating bi-allelic markers down gene trees remains the same as that employed in [7]. When requiring the data set to contain only polymorphic sites, if the generated site is not polymorphic, we discard both gene tree and markers, and repeat until a polymorphic site is generated.

## Results

### Simulations

#### The method’s ability to infer the phylogenetic network topology

We used the following commands in PhyloNet to generate 200 data sets, 20 replicates for each of the two model phylogenetic networks in Fig. 3 and each number of sites (*numsites* ∈ {100,1000,10000,100000,1000000)}:

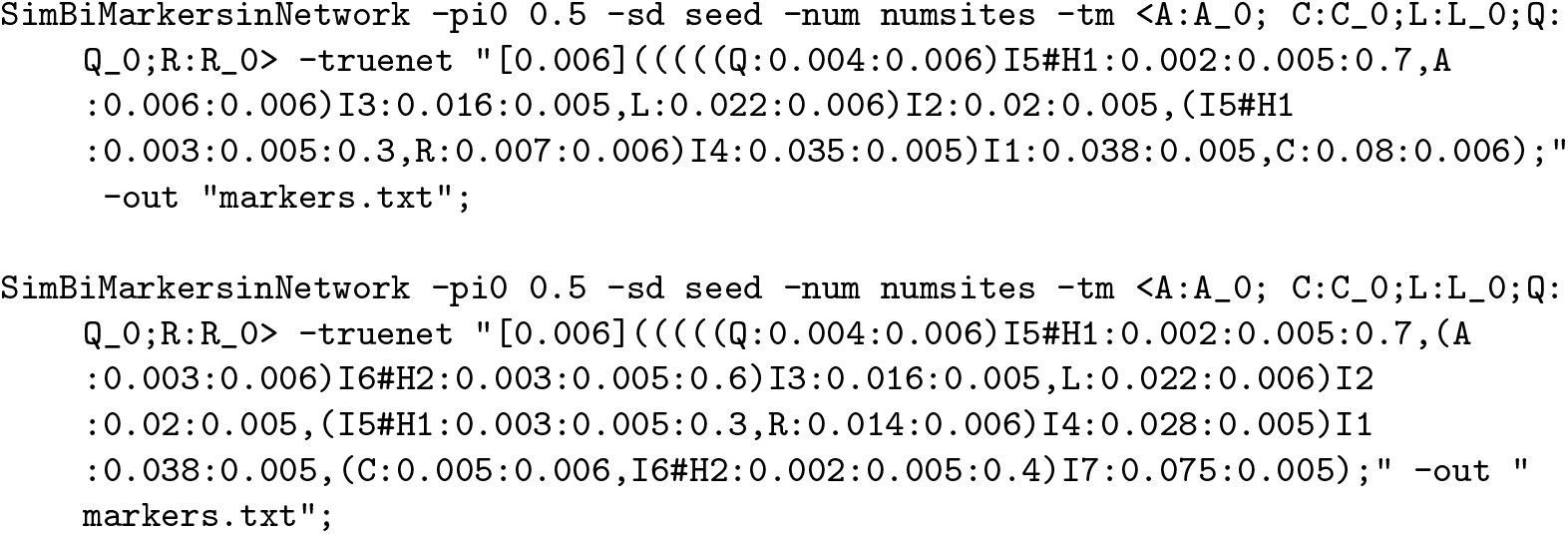

For the value of *seed* in the “-sd” option, we used a different 8-digit integer for each of the 20 replicates.

The true networks of those commands correspond to two models, given by the two phylogenetic networks, their branch lengths, and inheritance probabilities, shown in Fig. 3. These networks and parameters were inspired by the phylogenetic networks inferred from an empirical genomic data set in [21, 22].

**Fig 3.**
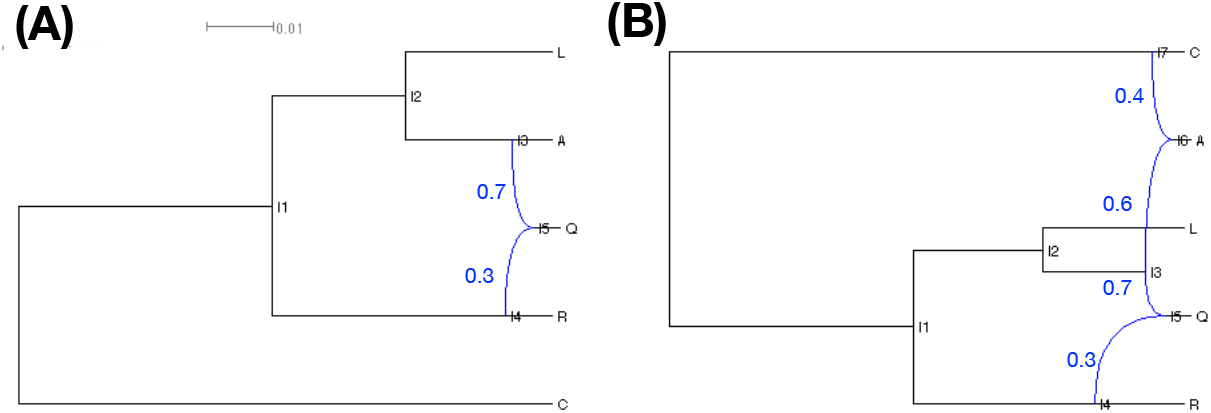
The two model phylogenetic networks used to generate the simulated data sets. The branch lengths of the phylogenetic networks are measured in units of expected number of mutations per site (scale is shown). The inheritance probabilities are marked in blue. Both networks are based on the same “backbone” tree: Removing the R→Q reticulation edge in (A) and the C→A and R→Q reticulation edges in (B) gives rise to the tree (C,(R,(L,(A,Q)))). The hybridization events in the panel can be viewed as involving pairs of branches of this tree: (A) The hybridization is from R to Q. (B) One hybridization is from R to Q and another is from C to A.

For each of the two models, we simulated data sets consisting of 100, 1000, 10000, 100000, and 1000000 bi-allelic sites, with one haploid generated for every taxon. In the simulations, we set *u* =1 and *υ* = 1 as the mutation rates. Furthermore, similar to [7], we used *θ* = 0.006 as the population mutation rate for external branches, and *θ* = 0.005 for internal branches and root, both in the unit of population mutation rate per site. Under these settings, we observed that each of the 200 data sets contained between 19% and 21% polymorphic sites; the remaining sites were all monomorphic.

To test the ability of our algorithm to recover the topology of the true phylogenetic network, we ran the RJMCMC sampler on the simulated data sets. To test how robust the method is to the setting of the prior on the population mutation rate, we ran the sampler under both the “correct” (*α* = 2, *β* = 0.003) and “incorrect” (*α* = 2, *β* = 0.0003) prior settings as in the following commands:

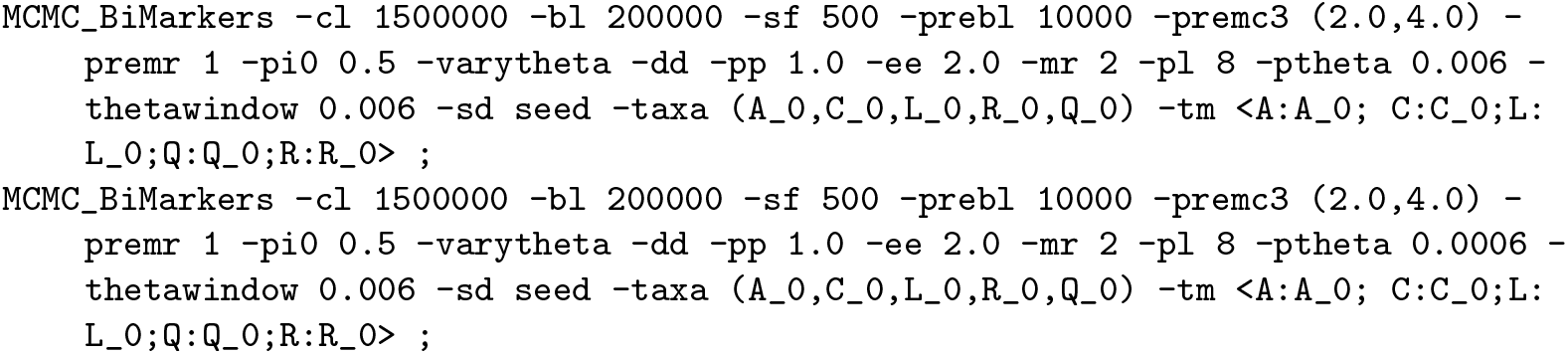

To speed up convergence, we used a pre-burn-in phase before we ran the actual sampling process. The pre-burn-in phase consisted of 3 chains with temperatures 1.0, 2.0, and 4.0, respectively, and each with 10,000 iterations. Following that, the actual sampling phase consisted of an MCMC chain for 1.5 × 10^6^ iterations, with a burn-in period of 2 × 10^5^ iterations, and one sample was collected from every 500 iterations. We used the network with the highest likelihood from the pre-burn-in phase as the starting network for the actual sampling phase. A maximum number of reticulations in this pre-burn-in phase was set at 1 and 2 for the data sets simulated on the networks in Fig. 3(A) and Fig. 3(B), respectively. In the actual sampling phase, the maximum number of reticulations was set to 2 and 3, respectively.

We summarized the results based on 1,040,000 samples (20 replicates, 2 networks, 2 prior settings, 5 different numbers of sites, and 2600 samples per chain). Fig. 4 shows the method’s performance in terms of the number of reticulation it infers. In this figure, for each number of sites, each bar corresponds to the ratio of networks (out of 52,000) with the specified number of reticulations. As the results show, the method almost always infers a tree when the number of sites is 100, for both true networks and regardless of the prior setting on the population mutation rate. When 1000 sites are used, the method infers the correct number of reticulations in almost 75% of the samples in the case of the true 1-reticulation network, whereas the rest of the samples are almost all trees. However, when 10,000 sites or more are used, almost all samples have the correct number of reticulations, for both model networks and regardless of the prior setting on the population mutation rate. In the case of the true 2-reticulation network, when 1,000 sites are used, the method almost always infers a 1-reticulation network.

**Fig 4.**
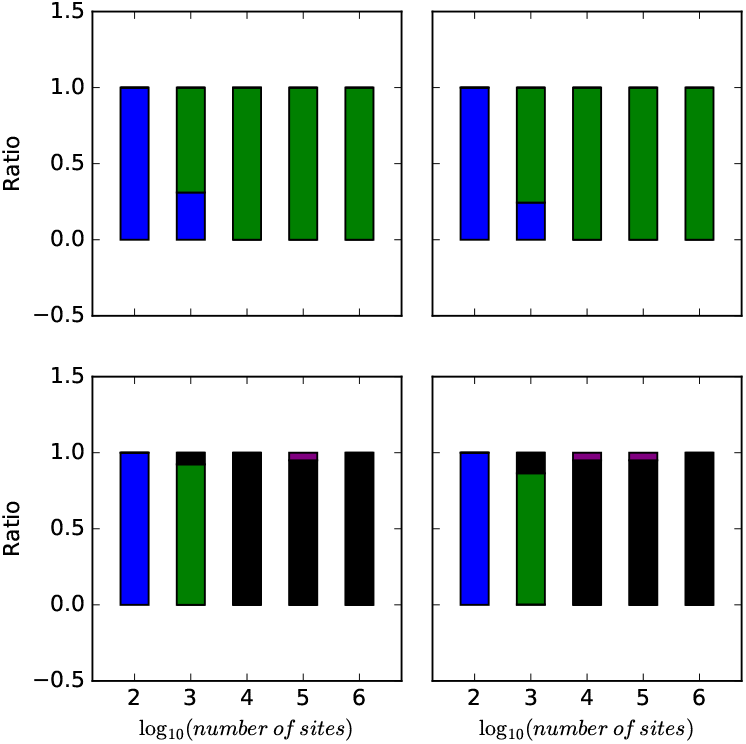
The ratio of trees (blue), 1-reticulation networks (green), 2-reticulation networks (black), and 3-reticulation networks (purple) sampled under different simulation settings. Top row: The true network is the 1-reticulation network in Fig. 3(A). Bottom row: The true network is the 2-reticulation network in Fig. 3(B). Left column: The correct prior hyperparameters for the population mutation rate were used. Right column: The incorrect prior hyperparameters for the population mutation rate were used.

While the method performs well in terms of estimating the number of reticulations, the next natural question is: Does the method infer the correct topology of the network? To answer this question, we assessed the topological differences between the sampled networks and the true ones in two different ways. When the method infers a network with at least one reticulation node, we compared the topology of the network to the true network topology using the dissimilarity measure of [35]. Furthermore, when the method infers a tree, we compared the tree to the “backbone tree” of the true network using the Robinson-Foulds metric [36]. Given a phylogenetic network with inheritance probabilities on its reticulation edges, removing for each reticulation node the incoming edge with the smaller inheritance probability results in a tree, which we call the backbone tree. For example, for the network in Fig. 3(B), the reticulation edges with inheritance probabilities 0.3 and 0.4 would be removed, resulting in the backbone tree (C,((L,(A,Q)),R)). It is important to note that in the presence of deep coalescence, it is extremely hard to capture the relationship between a phylogenetic network and its parental trees by a backbone tree [37]. The results for the simulated data sets are shown in Fig. 5.

**Fig 5.**
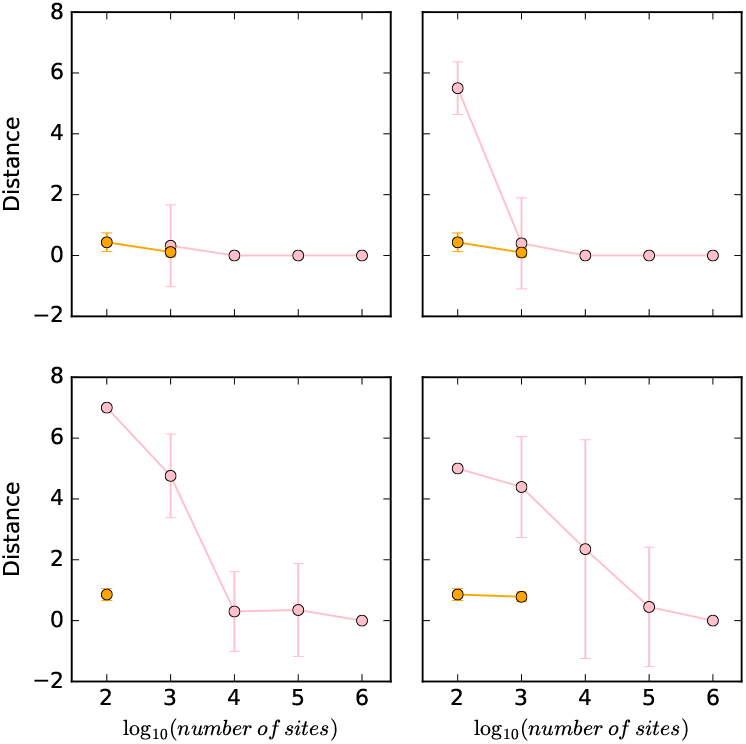
The topological distance (pink) between sampled networks and true network, and the Robinson-Foulds distance (orange) between sampled trees and true backbone tree, under different simulation settings. Top row: The true network is the 1-reticulation network in Fig. 3(A). Bottom row: The true network is the 2-reticulation network in Fig. 3(B). Left column: The correct prior hyperparameters for the population mutation rate were used. Right column: The incorrect prior hyperparameters for the population mutation rate were used.

In this figure, it is easy to think of the orange points as pertaining to the samples that correspond to the blue bars in Fig. 4, whereas the pink points pertain to all the other samples. Considering first the data sets simulated on the 1-reticulation network, the result indicate a very good performance. When a network is sampled, it is almost always the true network, as reflected by the topological distance close to 0. The only exception is the network inferred when using the incorrect prior hyperparameters on 100 sites. However, it is important to note that in most cases under this setting, a tree was sampled. Furthermore, when a tree was sampled by our method, its distance to the backbone tree is almost 0. In other words, when our methods failed to infer the true network, its because of lack of signal to infer the reticulation event, but it still inferred the underlying backbone tree.

In the case of the data sets simulated on the 2-reticulation network, the results differ. When the correct prior hyperparameters are used, 10,000 sites or more are needed to infer the true network. When the incorrect prior hyperparameters were used, the true network was sampled almost exclusively when 1,000,000 sites were used. When the method sampled a tree, that tree did not exactly match the backbone tree in this case, but was very close to it.

These results combined demonstrate a natural aspect of network inference: The more reticulate the evolutionary history, the more sites are needed for accurate inference of the network’s topology. In general, when the method makes a wrong inference, it is mostly in inferring the reticulation events themselves. However, in some cases, the tree inferred by the method does not match exactly the backbone tree, which is not unexpected given that the network in many cases is more than the sum of a backbone tree and a set of reticulation edges [37]. It is worth noting that while not many empirical AFLP-or SNP-based studies currently include as many as 10,000 loci, such large data sets may become commonplace as genomic technologies continue to advance.

#### The method’s ability to estimate the continuous parameters

We used the following command in PhyloNet [30] to generate 80 data sets on the network of Fig. 3(A) to test the ability of our algorithm to estimate the continuous parameters (branch lengths, inheritance probabilities, and population mutation rates):

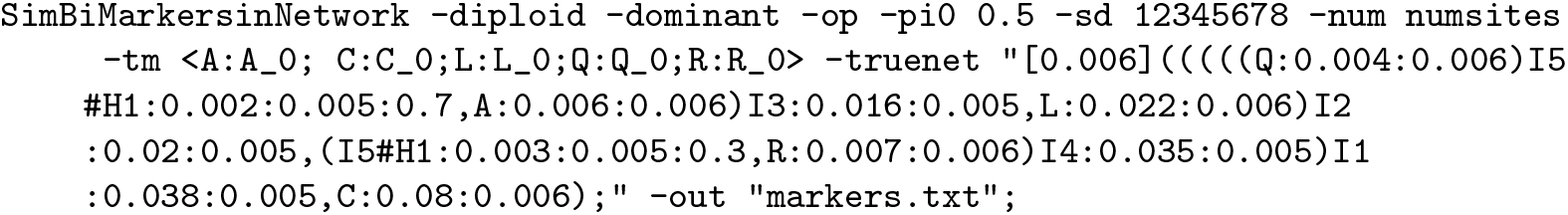

We used *numsites* ∈ {1000,10000,100000,1000000} given that the method almost never inferred a network when using only 100 sites. As before, we varied the value of *seed* for each of the 20 replicates. Here, we assumed one diploid genome per taxon, with dominant markers.

For the analysis, we first used a pre-burn-in phase with the exact same setting as above, and then started an RJMCMC chain from the network with the highest likelihood, using the following command:

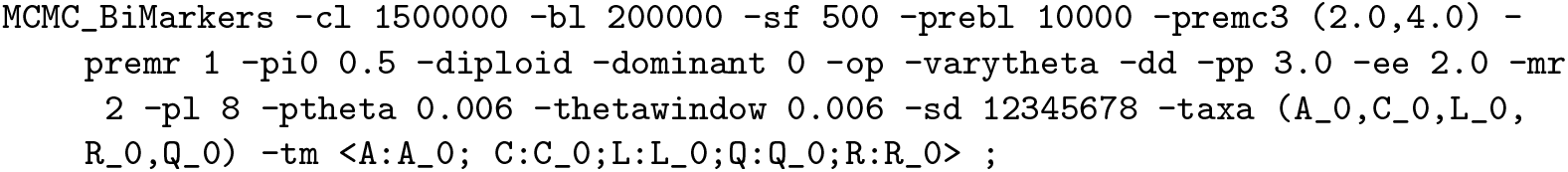

We ran the chain for 1.5 × 10^6^ iterations, with 2 × 10^5^ burn-in iterations, and one sample was collected from every 500 iterations. We summarized the results based on the collected samples whose topologies were the true network topology (otherwise, there is no easy way to find correspondence between the parameters in the inferred networks and those in the true one) in the form of histograms.

Histograms of the sampled branch lengths are shown in Fig. 6. The results indicate a very good performance of the method. The histograms peak around the true value for all branches, regardless of the number of sites used. However, as the number of sites increases, the variance around the true value shrinks.

**Fig 6.**
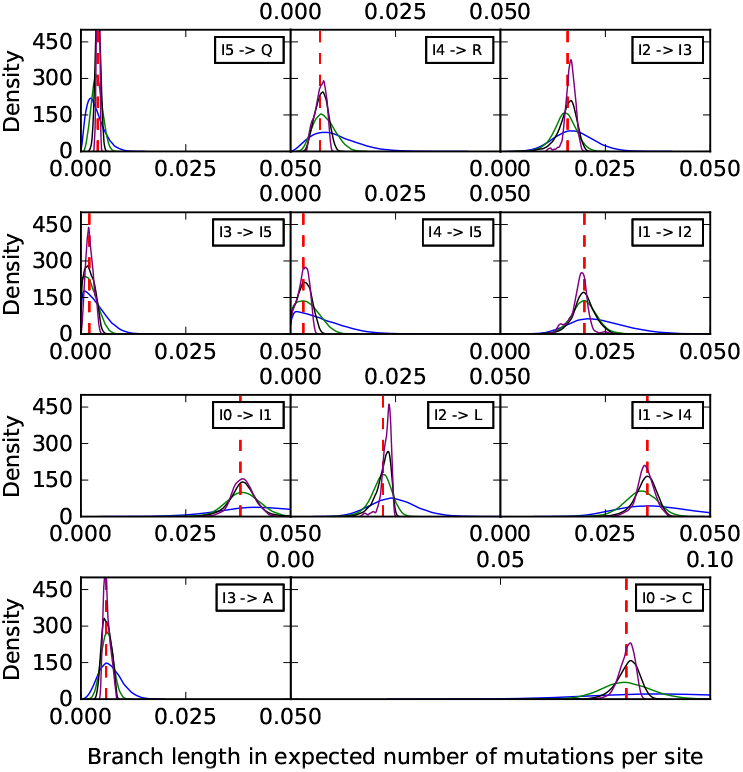
Histograms of the branch lengths sampled by our method on the simulated data set corresponding to the phylogenetic network of Fig. 3(A). Blue: 1,000 sites. Green: 10,000 sites. Black: 100,000 sites. Purple: 1,000,000 sites. The red dashed lines correspond to the true values.

Histograms of the sampled population mutation rates are shown in Fig. 7. Unlike the case of branch lengths, we observe here that the population mutation rates of the external branches are well estimated, with the distribution tightening around the true values as the number of sites increases. However, the population mutation rates seem to be unidentifiable by the method for internal branches, especially those closer to the root. As we show below, increasing the number of individuals sampled per species help improve these estimates.

**Fig 7.**
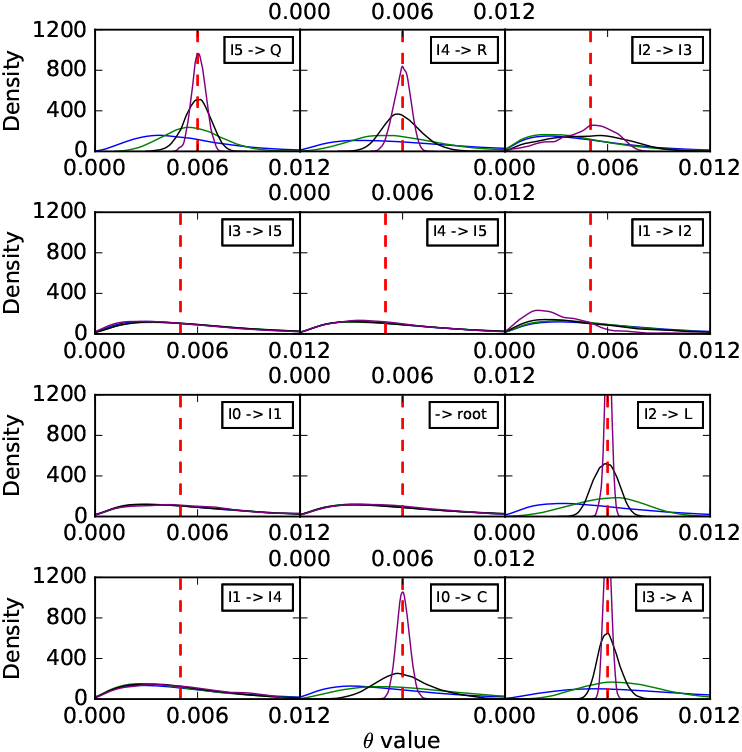
Histograms of the population mutation rates sampled by our method for each of the branches on the simulated data set corresponding to the phylogenetic network of Fig. 3(A). Blue: 1,000 sites. Green: 10,000 sites. Black: 100,000 sites. Purple: 1,000,000 sites. The red dashed lines correspond to the true values.

Finally, a histogram of the sampled inheritance probabilities is shown in Fig. 8. The method does a very good job at sampling the true value in this case. As the number of sites increases, the distribution of sampled value tightens around the true value and becomes almost all concentrated within 0.1 around it when 1,000,000 sites are used.

**Fig 8.**
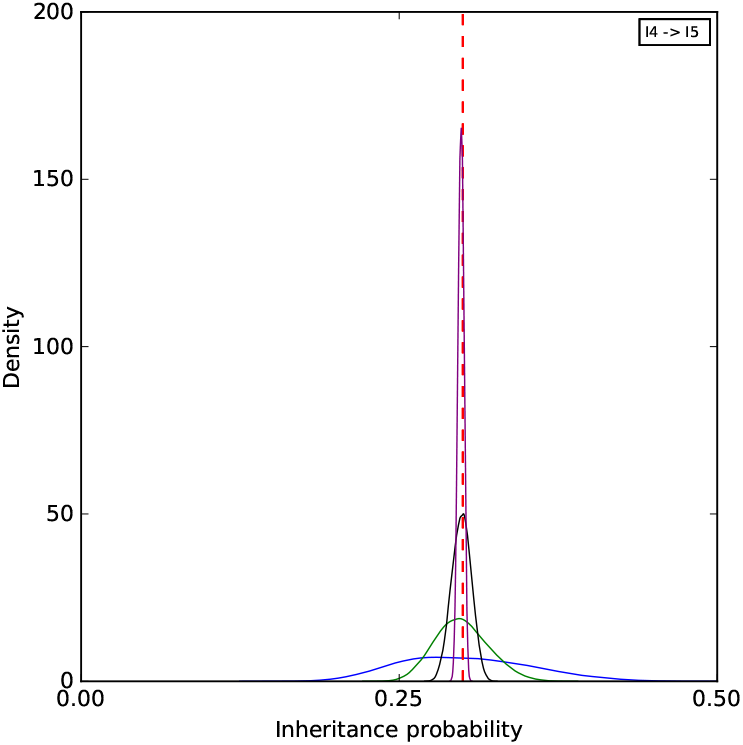
A histogram of the inheritance probabilities sampled by our method on the simulated data set corresponding to the phylogenetic network of Fig. 3(A). Blue: 1,000 sites. Green: 10,000 sites. Black: 100,000 sites. Purple: 1,000,000 sites. The red dashed line corresponds to the true values.

To further assess the mixing of our sampler, we also performed five independent runs on an arbitrary data set with 10,000 sites, where each run consisting of an MCMC chain under the same settings as above. The results for the continuous parameters are shown in Figs. 9–11. Samples from the different runs are in very good agreement, further indicating good mixing.

**Fig 9.**
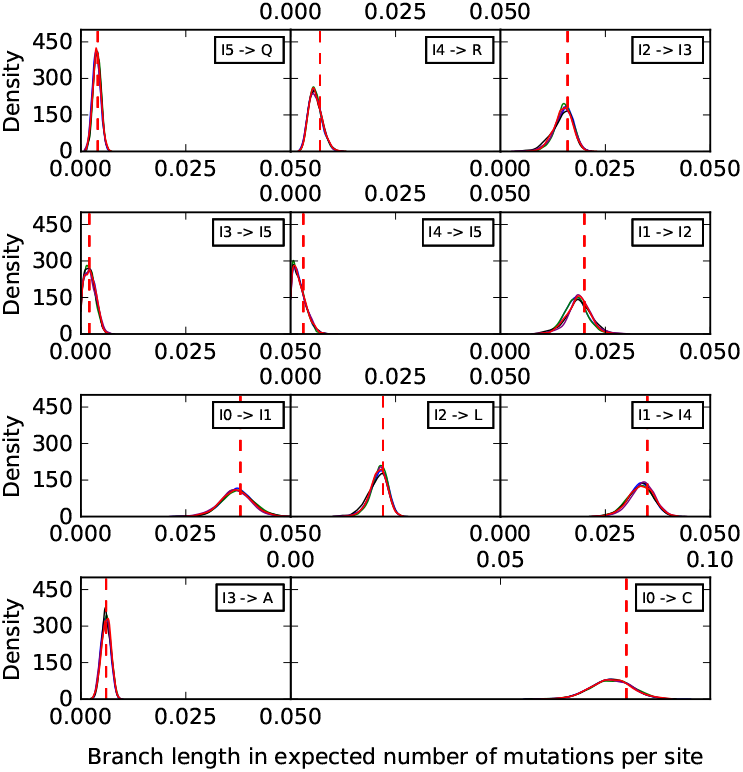
Histograms of the branch lengths sampled by our method on the simulated data set corresponding to the phylogenetic network of Fig. 3(A). The five curves correspond to five independent runs. The red dashed lines correspond to the true values.

#### The effect of the number of sampled individuals on parameter estimates

Monomorphic sites help estimate parameter values, but sometimes they are removed because they are uninformative for estimating the topology and to reduce the computation time for the phylogenetic analyses. If there are only polymorphic sites in the data set, sampling multiple individuals could improve parameter estimation. To investigate this aspect, we set up a simulation with the phylogenetic network in Fig. 12 (we reduced the number of taxa to make the running time over many data sets manageable). In the simulation, we set *u* =1 and *υ* = 1 as the mutation rates. Furthermore, we used *θ* = 0.005. We sampled one diploid individual for each of the three species A, B, and D, and four diploid individuals for species C. We generated 10,000 polymorphic sites with dominant markers for each of those individuals, with 5 replicates.

We used following command in PhyloNet to generate the data set:

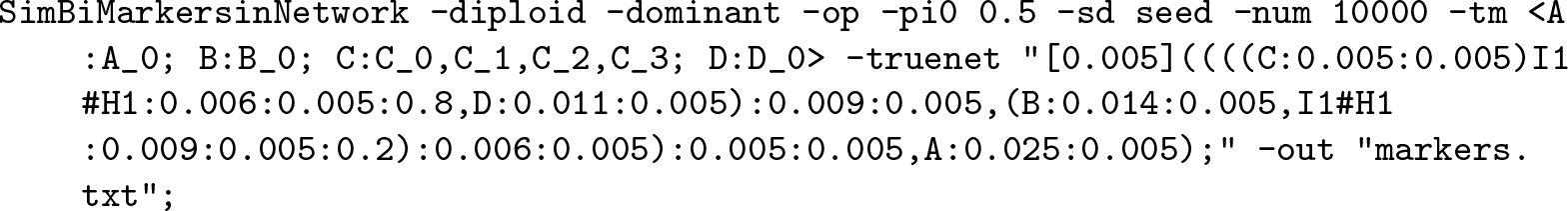

**Fig 10.**
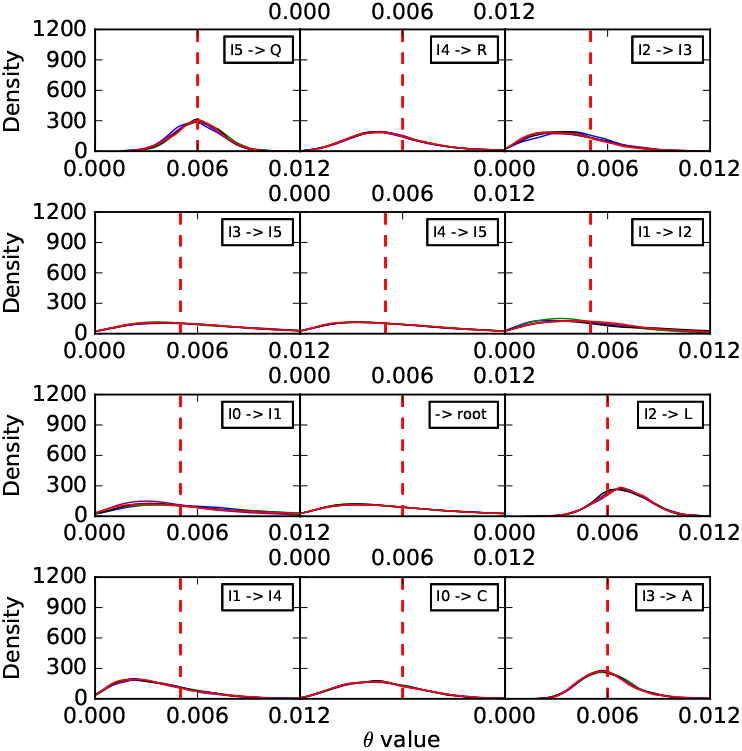
Histograms of the population mutation rates sampled by our method for each of the branches on the simulated data set corresponding to the phylogenetic network of Fig. 3(A). The five curves correspond to five independent runs. The red dashed lines correspond to the true values.

We ran the method on the entire data set (7 diploid individuals, amounting to 14 haploid individuals), and on a subset that consists of a single diploid individual from each of the four species (8 haploids in total).

We ran each MCMC chain for 5 × 10^5^ iterations, with 5 × 10^4^ burn-in iterations, and one sample was collected from every 500 iterations. We used the collected samples whose topologies were the true network topology to summarize the results in the form of histograms of the parameter estimates.

Histograms of the sampled branch lengths are shown in Fig. 13. As the results show, the inclusion of multiple individuals from the hybrid species (C) helps most in estimating the lengths of the three branches surrounding the reticulation event (the three edges incident with node I1).

**Fig 11.**
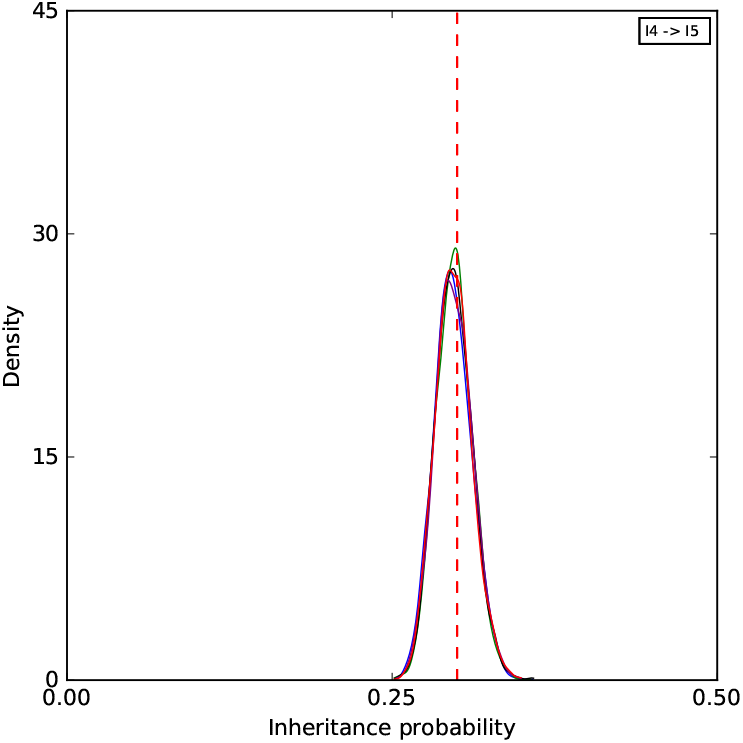
A histogram of the inheritance probabilities sampled by our method on the simulated data set corresponding to the phylogenetic network of Fig. 3(A). The five curves correspond to five independent runs. The red dashed line corresponds to the true value.

**Fig 12.**
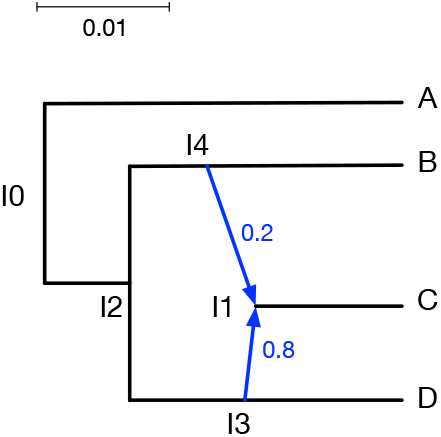
The phylogenetic network used to investigate effect of multiple individuals. The branch lengths of the phylogenetic networks are measured in units of expected number of mutations per site. The inheritance probabilities are marked in blue.

Histograms of the sampled population mutation rates are shown in Fig. 14. In this case, the most improvement gained from the inclusion of multiple individuals pertains to the external branch to taxon C.

A histogram of the sampled inheritance probabilities is shown in Fig. 15. The inheritance probability estimates improve when four individuals are sampled, as the distribution of the sampled values become more concentrated and peak much closer to the true value.

This simulation was performed on DAVinCI, which is a batch scheduled High-Throughput Computing (HTC) cluster. We used 6 cores, with one thread per core running at 2.83GHz, and 4G RAM per thread. The average runtime for analyzing the full data set with four individuals sampled from C is 47.5 hours for each replicate. The average runtime for analyzing the subset with a single individual sampled from C is 0.3 hour for each replicate. This shows the drastic effect of the number of individuals sampled on the running time of the method.

**Fig 13.**
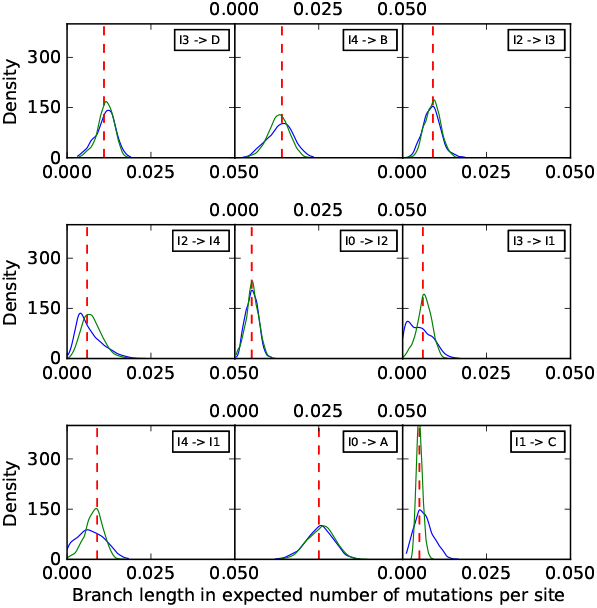
Histograms of the branch lengths sampled by our method on the simulated data set corresponding to the phylogenetic network of Fig. 12. In all cases, a single diploid individual was sampled from A, B, and D. Blue: A single diploid individual is sampled from C. Green: Four diploid individuals are sampled from C. The red dashed lines correspond to the true values.

**Fig 14.**
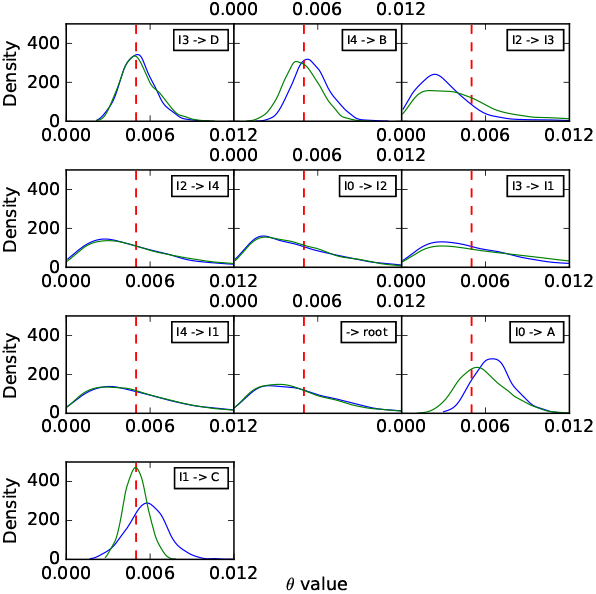
Histograms of the population mutation rates sampled by our method for each of the branches on the simulated data set corresponding to the phylogenetic network of Fig. 12. In all cases, a single diploid individual was sampled from A, B, and D. Blue: A single diploid individual is sampled from C. Green: Four diploid individuals are sampled from C. The red dashed lines correspond to the true values.

#### The method’s robustness to violations in the assumptions

To study the robustness of our method to violations in the underlying assumptions of the model, we simulated data sets on the network of Fig. 3(A) with 100, 200, 300, 400, and 500 sites under different conditions.

Our model assumes the markers are unlinked. To simulate data with linked markers, we used the following command to produce multiple markers coming from the same gene tree:

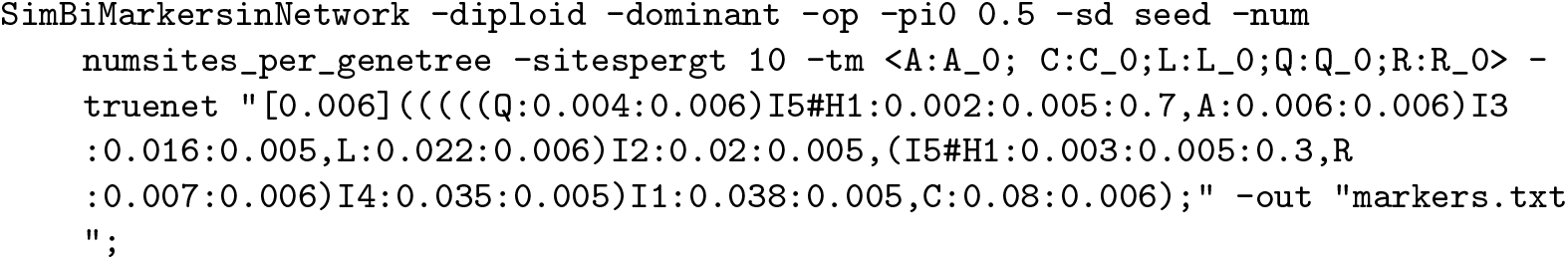

**Fig 15.**
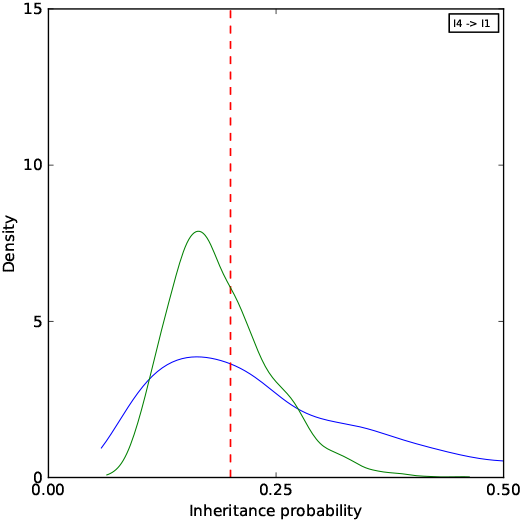
A histogram of the inheritance probabilities sampled by our method on the simulated data set corresponding to the phylogenetic network of Fig. 12. In all cases, a single diploid individual was sampled from A, B, and D. Blue: A single diploid individual is sampled from C. Green: Four diploid individuals are sampled from C. The red dashed lines correspond to the true value.

Here, *numsites*_*per*_*genetree* was once set to 10 to generate 10 markers per gene tree, and then set to 100 to generate 100 markers per gene tree. Naturally, the number of gene trees generated was divided by either 10 or 100, respectively. Each setting was used to generate 100, 200, 300, 400, 500 sites in total, and with 5 replicates using different integers as the random seeds.

Our model assumes a constant rate of mutation across all lineages and all markers. We simulated data sets with rate variation across markers and lineages. In the second step of our simulator, we adopted a rate variation which mimics the *GT R* + *I* + Γ setting in [38]. The following is one of the commands we used to simulate rate variation:

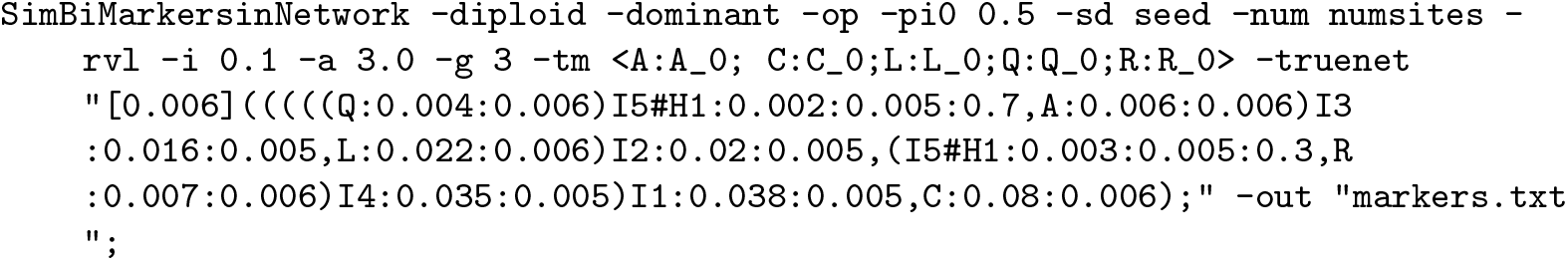

The command was repeated with “-rvl -i 0.1 -a 3.0 -g 3”, “-rvl -i 0.2 -a 5.0 -g 3”, “-rvm -i 0.1 -a 3.0 -g 3”, “-rvm -i 0.2 -a 5.0 -g 3”. Here, “-rvl” means rate variation across lineages, “-rvm” means rate variation across markers, “-i” indicates the proportion of invariable sites, “-a” indicates the shape for the gamma rate heterogeneity, and “-g” indicates the number of categories for the discrete gamma rate heterogeneity model. Five replicates were generated for each setting, using different integers as the random seeds.

Furthermore, in our Bayesian formulation given by Eq. (16), the prior *p*(Ψ) includes a Poisson distribution on the number of reticulation nodes in the network. In our experiments here, we varied the mean of the Poisson distribution in {1, 3}.

In total, there are 8 settings. For each setting, we generated five replicate 100-, 200-, 300-, 400-, 500-site data sets.

As above, we employed a pre-burn-in phase to obtain a good starting point, from which we then started an MCMC chain for each data set with 1.5 × 10^6^ iterations, 2 × 10^5^ burn-in iterations, and one sample collected from every 500 iterations. In total, there were 13,000 samples collected for each data set with different number of sites and different settings.

To avoid reporting results for all branch lengths in the sampled networks, Fig. 16 shows the estimation of the height of trees and networks sampled in each setting. The five bars from bottom to top correspond to the minimum, first-, second-, third-quantile, and the maximum. The most important striking pattern in the figure is that as the number of sites increases, the estimate of the height converges to the true value and the variance in the estimated value becomes smaller. In all cases of assumption violations, 500 sites are sufficient to obtain a very accurate estimate of the height.

**Fig 16.**
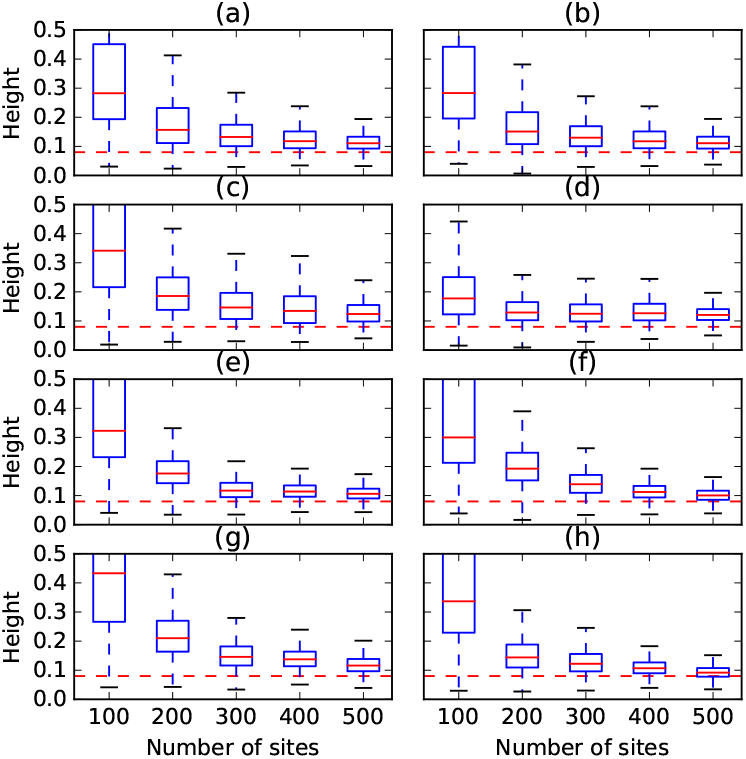
The height of trees and networks sampled under different simulation settings and violations in the different assumptions. The red dashed lines correspond to the true values. In each panel at most one condition is violated. (a) Mean of 1.0 is used for the Poisson prior on the number of reticulations. (b) Mean of 3.0 is used for the Poisson prior on the number of reticulations. (c) Linked loci: 10 sites are generated per gene tree. (d) Linked loci: 100 sites are generated per gene tree. (e) Rate variation across lineages with 0.1 of invariable sites and 3.0 as shape of gamma rate heterogeneity. (f) Rate variation across lineages with 0.2 of invariable sites and 5.0 as shape of gamma rate heterogeneity. (g) Rate variation across markers with 0.1 of invariable sites and 3.0 as shape of gamma rate heterogeneity. (h) Rate variation across markers with 0.2 of invariable sites and 5.0 as shape of gamma rate heterogeneity.

Fig. 17 shows the estimates of the number of reticulations under the different settings and violations. In all cases, 500 sites were sufficient to recover the true number of reticulations, regardless of the violation in assumptions. Interestingly, the set mean of the Poisson distribution on the number of reticulations has an almost identical effect regardless of whether it is set to 1 . 0 or to 3. 0. The inclusion of linked loci has much more of an effect on the number of reticulations inferred. As can be seen in the figure, even with 400 sites, the method still infers trees in many of the samples. Rate variation across lineages and sites has a similar effect, which is overcome by the method when 400 sites or more are used.

The topological differences between the inferred networks and true one and between the inferred trees and the true backbone tree are shown in Fig. 18. With the exception of the 100-site data sets, the method is very robust in terms of the topology of the inferred network. When it infers a network, it almost always obtains the true one. When it infers a tree, it almost always obtains the backbone tree.

Put together, these results point to a very robust method. In particular, when at least 500 sites are used, the method is almost completely robust to any of the violations we studied here.

**Fig 17.**
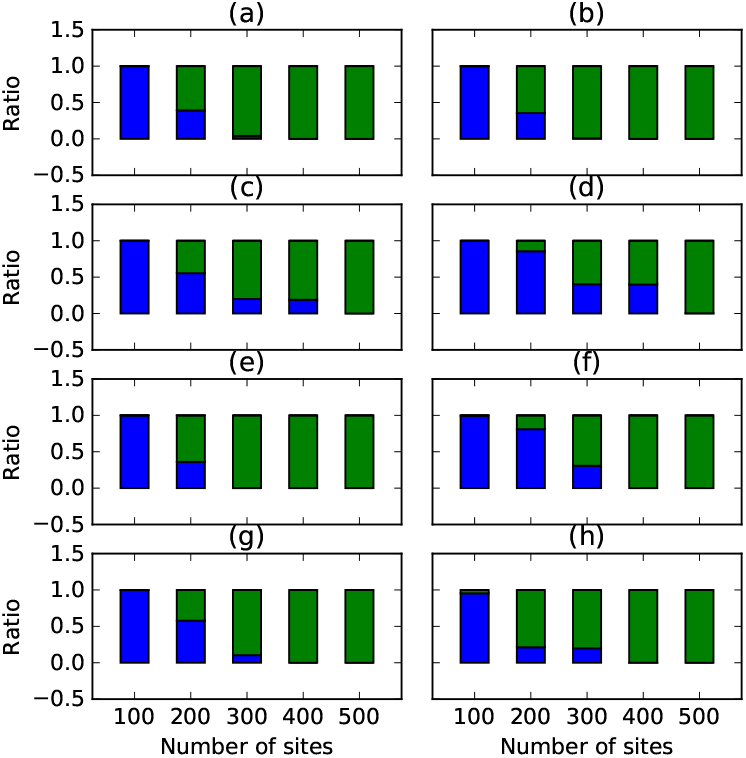
The ratio of trees (blue) and 1-reticulation networks (green) sampled under different simulation settings and violation in the different assumptions. The true number of reticulations is 1. In each panel at most one condition is violated, (a) Mean of 1.0 is used for the Poisson prior on the number of reticulations, (b) Mean of 3.0 is used for the Poisson prior on the number of reticulations, (c) Linked loci: 10 sites are generated per gene tree, (d) Linked loci: 100 sites are generated per gene tree, (e) Rate variation across lineages with 0.1 of invariable sites and 3.0 as shape of gamma rate heterogeneity, (f) Rate variation across lineages with 0.2 of invariable sites and 5.0 as shape of gamma rate heterogeneity, (g) Rate variation across markers with 0.1 of invariable sites and 3.0 as shape of gamma rate heterogeneity, (h) Rate variation across markers with 0.2 of invariable sites and 5.0 as shape of gamma rate heterogeneity.

### Analysis of an empirical data set

Two small subsets of a larger AFLP data set of multiple New Zealand species of the plant genus *Ourisia* (Plantaginaceae) [39] were analyzed, including previously unpublished AFLP profiles from two different hybrid individuals *O.* × *cockayneana* and *O.* × *prorepens* (herbarium codes follow [40] [continuously updated]). There are both morphological [41] and molecular (Meudt unpubl.) data supporting the hybrid nature of these two individuals. Although other *Ourisia* hybrid combinations have been reported in New Zealand [41], *O.* × *cockayneana* and *O.* × *prorepens* are perhaps the most common, both involve *O. caespitosa* as a putative parent, and both have been formally named. Each data subset comprised five diploid individuals in total, which means ten haploid individuals were effectively analyzed due to the correction for dominant markers.

A Poisson distribution with λ = 1.5 as the prior on the number of reticulations, an exponential prior with λ = 2.0 as the prior on the species divergence times, and a Gamma distribution with *α* = 2.0 and *β* = 0.05 as the prior on the population mutation rates were adopted. An MCMC chain was run on each data subset for 1.5 × 10^6^ iterations with 2 × 10^5^ burn-in iterations, and a sample was collected every 500 iterations. We used following commands:

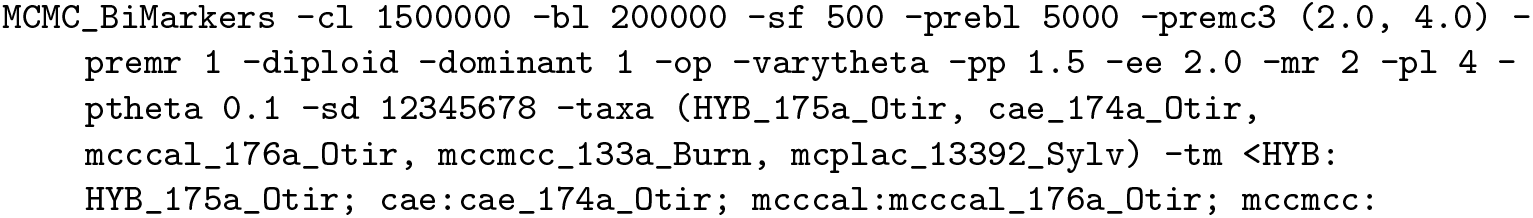

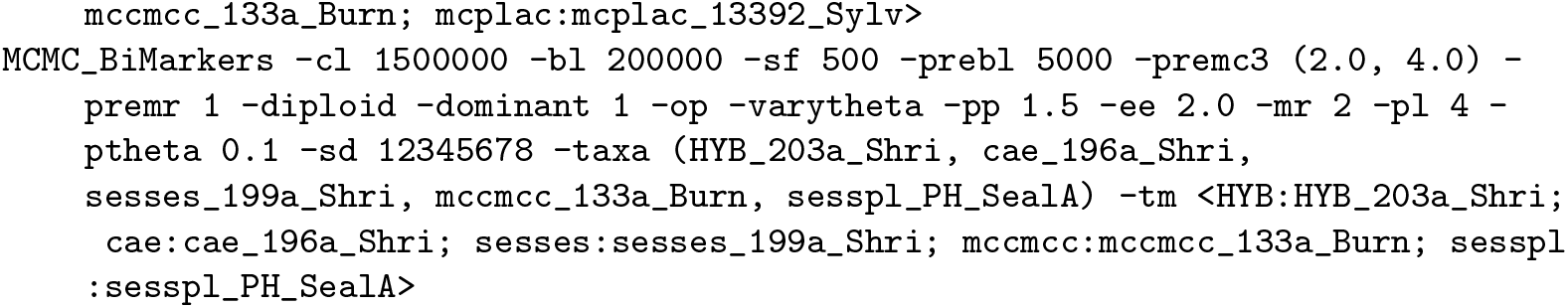

**Fig 18.**
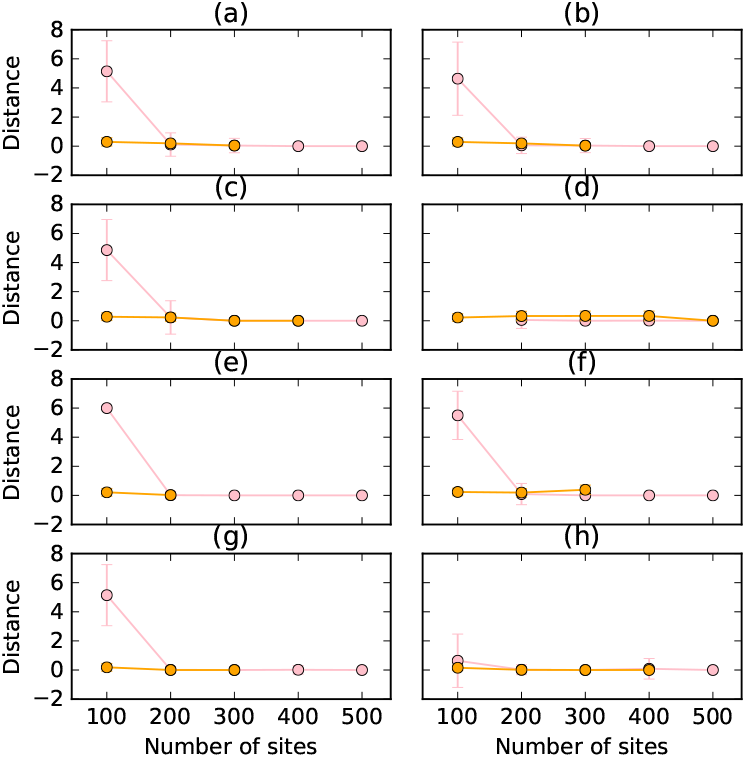
The topological distance (pink) between sampled networks and true network, and the Robinson-Foulds distance (orange) between sampled trees and true backbone tree, under different simulation settings and violation in the different assumptions. In each panel at most one condition is violated, (a) Mean of 1.0 is used for the Poisson prior on the number of reticulations, (b) Mean of 3.0 is used for the Poisson prior on the number of reticulations, (c) Linked loci: 10 sites are generated per gene tree, (d) Linked loci: 100 sites are generated per gene tree, (e) Rate variation across lineages with 0.1 of invariable sites and 3.0 as shape of gamma rate heterogeneity, (f) Rate variation across lineages with 0.2 of invariable sites and 5.0 as shape of gamma rate heterogeneity, (g) Rate variation across markers with 0.1 of invariable sites and 3.0 as shape of gamma rate heterogeneity, (h) Rate variation across markers with 0.2 of invariable sites and 5.0 as shape of gamma rate heterogeneity.

#### Data subset with hybrid *O.* × *cockayneana*

The first data subset comprises the following five individuals: *O. macrocarpa* (voucher: *Meudt 133a,* MPN 29546; herbarium codes follow [40] [continuously updated]), *O. macrophylla* subsp. *lactea* (*Cameron 13392,* AK 294893), hybrid *O.* × *cockayneana* (*Meudt 175a,* MPN 29710), *O. caespitosa* (*Meudt 174a,* MPN 29705), and *O. calycma* (*Meudt 176a,* MPN 29713). The number of loci in this data set is 802.

The maximum a posteriori (MAP) phylogenetic network is shown in Fig. 19. The effective sample size was larger than 2,000. All topologies sampled successfully detected the hybrid and its putative parents. If the hybrid is removed, the topology in Fig. 19 also agrees with that of Fig. 3 in [39].

It should be noted that the posterior standard deviations reported in Fig. 19 are much larger than those in [7]. This is perhaps not unexpected because we only used one individual per species in our analysis. Our simulation study shows that increased sampling of individuals helps the estimation of parameters, whereas when only one individual per species is sampled, the posterior distribution is much wider.

**Fig 19.**
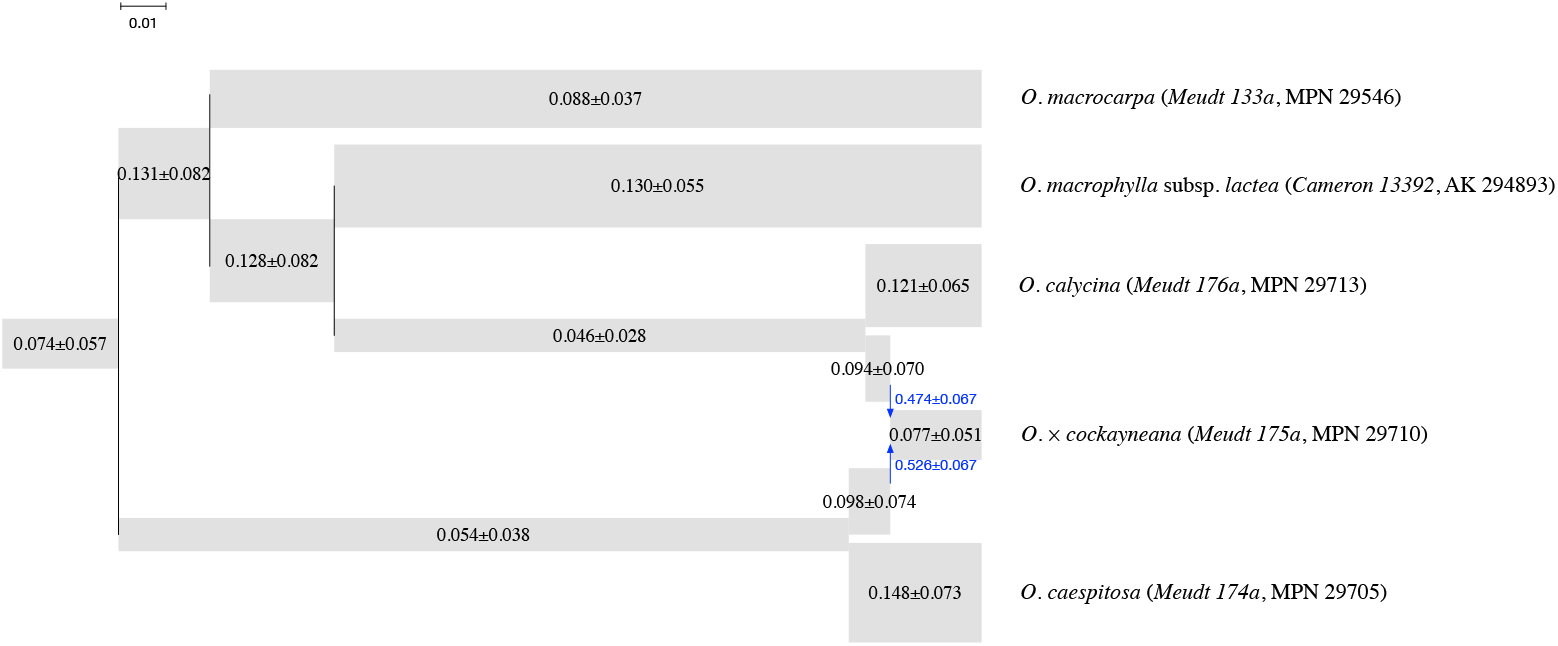
The MAP phylogenetic network for the subset with the hybrid *O.* × *cockayneana* (*Meudt 175a*, MPN 29710) and putative parents. The width of each tube is proportional to the population mutation rate of each branch, which is printed on each tube. The length of each tube is proportional to the length of the corresponding branch in units of expected number of mutations per site (scale shown). Blue arrows indicate the reticulation edges and their inheritance probabilities are printed in blue.

#### Data subset with hybrid *O*. × *prorepens*

The second data subset comprises *O*. *sessilifolia* subsp. *splendida* (*Heenan s.n.,* MPN 32149), *O. macrocarpa* (*Meudt 133a,* MPN 29713), hybrid *O*. × *prorepens* (*Meudt 203^<u>a</u>^*, MPN 29774), *O. sessilifolia* subsp. sessilifolia (*Meudt 199a,* MPN 29771), and *O. caespitosa* (*Meudt 196a,* MPN 297695). The number of loci in this data set is 820.

The MAP phylogenetic network is shown in Fig. 20. The effective sample size was also larger than 2,000 in this case. The result shows our method successfully detected the hybrid and its putative parents. If the hybrid is removed, the topology in Fig. 20 also agrees with that of Fig. 3 in [39]. As with the first data subset, the posterior standard deviations reported in Fig. 20 are large.

Nevertheless, the mean values of inferred parameters are very similar for the two species that were common to the two data subsets, *O. caespitosa* and *O. macrocarpa.* The mean value of inferred population mutation rate of their corresponding leaves are similar. This shows that the method is both robust and consistent.

In summary, our method was able to extract the signal of the hybrid and successfully recover its putative parents, as well as reconstruct network topologies which were consistent with a previous study of a larger dataset [39].

## Discussion

Phylogenetic networks allow for representing evolutionary relationships that involve both vertical and horizontal transmission of genetic material. Extensions of the multispecies coalescent process to include hybridization events have facilitated the development of statistical methods for inferring and analyzing phylogenetic networks from gene tree estimates and sequence data. A major challenge with using gene tree estimates as the input to species phylogeny inference methods is the error in these estimates. While using the sequence data directly overcomes this issue, the problem of recombinations within loci can confound inferences. Using bi-allelic markers from individual, independent loci could provide a way to avoid both the gene tree uncertainty and recombination problems (the two are not necessarily independent). Furthermore, it is important to note that many biological studies use data sets that consists of bi-allelic markers and no available sequence alignment data for individual loci.

**Fig 20.**
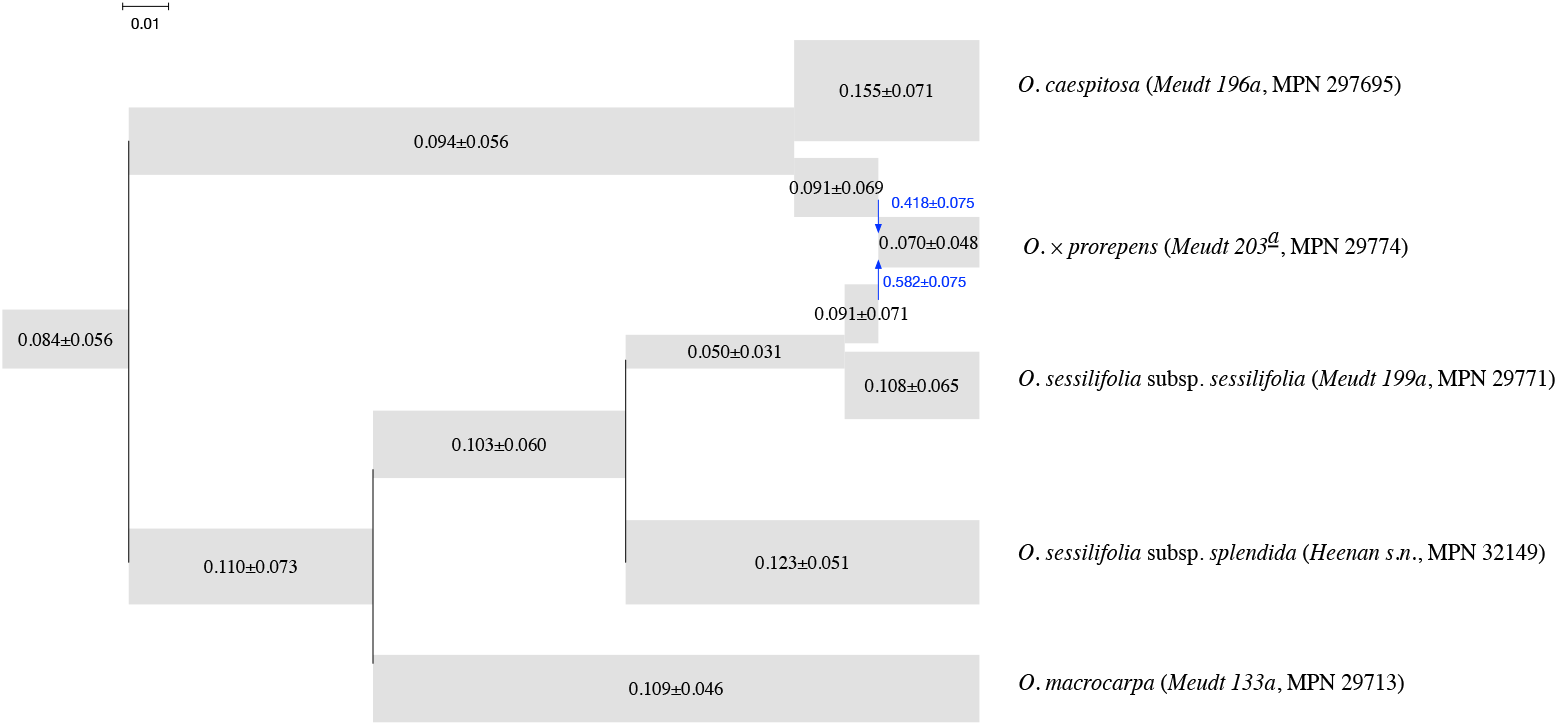
The MAP phylogenetic network for the subset with the hybrid *O*. × *prorepens* (*Meudt 203^<u>a</u>^*, MPN 29774) and putative parents. The width of each tube is proportional to the population mutation rate of each branch, which is printed on each tube. The length of each tube is proportional to the length of the corresponding branch in units of expected number of mutations per site (scale shown). Blue arrows indicate the reticulation edges and their inheritance probabilities are printed in blue.

Bryant *et al.* recently devised an algorithm for inferring species trees from bi-allelic genetic markers while analytically integrating out the gene trees for the individual loci [7]. In this paper, we extended their algorithm significantly so as the likelihood of a phylogenetic network given bi-allelic markers could be computed while integrating out the gene trees. This method complements existing ones that use gene tree estimates or sequence alignments as input for statistical inference of phylogenetic networks.

We implemented a Bayesian method for sampling the posterior of phylogenetic networks and their associated parameters from bi-allelic data, and studied its performance on both simulated and empirical data. The results indicate a very good performance of the method. This work adds a powerful method to the biologist’s toolbox that allows for estimating reticulate evolutionary histories.

A major bottleneck of the method is its computational requirements. While the SNAPP method is very time consuming on species trees, our method is much more time consuming given that reticulations in the phylogenetic network give rise to an explosion of the number of partial likelihoods that need to be computed and stored. More generally, the number of taxa in a data set has more of an effect on the running time of the method than the number of loci does. In particular, two aspects of the phylogenetic network under consideration affect the computational requirements of the method: The number of leaves under the reticulation nodes and the diameter of each of the reticulation nodes. As discussed above, the set of lineages entering a reticulation node must be bipartitioned in every possible way. This number of lineages is dependent on the number of leaves under that reticulation node. For example, if a single individual is sampled from a single species that exist under the reticulation node, then the number of bipartitions is very small (only two bipartitions exist). However, if *n* individuals are sampled from a single species that exist under the reticulation node or one individual is sampled per *n* species that exist under the reticulation node, then a number of bipartitions on the order of 2^*n*^ arises. This computation becomes much more demanding if there are more reticulation nodes on the path to a lowest articulation node. As for the diameter—which is the number of branches on the paths between the two parents of the reticulation node and a lowest articulation node above them, the larger its value, the more demanding the computation becomes. An important direction for future research is improving the computational requirements of the method to scale up to data sets with many taxa.

